# Bidirectional coupling among EMT, AXL-RB1 signaling and lineage switch drives resistance to osimertinib and worse clinical outcomes in NSCLC

**DOI:** 10.64898/2026.04.21.719547

**Authors:** Shreya Vashista, Ritesh Kumar Meena, Prakash Kulkarni, Ravi Salgia, Anish Thomas, Mohit Kumar Jolly

## Abstract

Acquired resistance to osimertinib remains a major barrier in EGFR-mutant lung adenocarcinoma (LUAD), and in many patients cannot be explained by secondary targetable mutations. This pattern highlights a central role for non-genetic plasticity programs, including epithelial–mesenchymal transition (EMT), drug tolerance, immune evasion, and lineage switch. Here, we used a systems-level framework to define how these processes are coordinated. We constructed a minimal gene regulatory network integrating core EMT regulators with AXL, RB1, PD-L1, and NF-κB, and analysed its emergent behaviour using dynamical simulations. The network resolved into two mutually inhibitory, self-reinforcing “teams”: an epithelial/sensitive team centred on RB1, miR-200, miR-34, p53, and E-cadherin, and a mesenchymal/resistant team centred on ZEB1, SNAIL, AXL, PD-L1, and NF-κB. Simulations predicted a strong coupling between EMT and osimertinib resistance, which was validated across bulk transcriptomic datasets from NSCLC cell lines, EGFR-mutant patient cohorts, and perturbation experiments. Inducing EMT increased RB1-loss programs, whereas osimertinib exposure induced AXL and EMT programs, supporting bidirectional regulation and reinforcement. Single-cell and spatial transcriptomic analyses further showed that EMT, AXL, PD-L1 activity, and reduced RB1 signaling co-occur within tumors. Clinically, activation of individual axes such as EMT, RB1 loss, or PD-L1 upregulation was associated with worse outcomes, while combined activation produced markedly poorer survival than any single axis alone. Extending the network to incorporate lineage regulators further linked a partial LUAD-to-LUSC shift with EMT, RB1 loss, and resistance. Together, these findings identify a network topology that coordinates multiple plasticity programs driving osimertinib resistance and suggest that disrupting this cooperative architecture may offer a therapeutic strategy in EGFR-mutant LUAD.

## Introduction

Lung cancer is the leading cause of cancer related mortality worldwide, with an estimated 1.8 million deaths, and low 5-year survival rate of 10-20% (1). Its two main histological types are non-small cell lung cancer (NSCLC) and small-cell lung cancer (SCLC). NSCLC accounts for 85% of lung cancer with lung adenocarcinoma (LUAD) being the most prevalent subtype (2). Mutations in epidermal growth factor receptor (EGFR) such as exon 19 deletion and exon 21 L858R substitutions are among the most well-characterized drivers of NSCLC, leading to constitutive activation of the receptor. EGFR tyrosine kinase inhibitors (EGFR-TKIs) are used to block EGFR activity, leading to improved clinical outcomes, but secondary mutations, T790M being the most prevalent one, led patients to develop resistance to first- and second-generation EGFR-TKIs such as erlotinib or gefitinib. Osimertinib, a third-generation EGFR-TKI, has redefined the standard of care for EGFR-mutant NSCLC, being effective against T790M-positive lung cancer (3). However, the emergence of acquired resistance to osimertinib limits its long-term efficacy and demands a better understanding of underlying mechanisms to improve the clinical management of NSCLC.

While some genomic drivers of osimertinib resistance have been reported, such as EGFR C797S mutation, approximately 50% of patients exhibiting resistance lack any identifiable mutations, emphasizing the role of non-genetic cell adaptation in evading drug response (4). Therapeutic options for these patients with no targetable mutations remain quite limited. Many EGFR-independent mechanisms such as an epithelial-mesenchymal transition (EMT) (5), drug-tolerant persisters (DTPs) (6) and lineage switch to SCLC (7) or lung squamous cell carcinoma (LUSC) (8) have been implicated in enabling resistance to osimertinib. The loss of tumour suppressor gene RB1 is observed frequently in EGFR mutant LUAD undergoing lineage plasticity to SCLC, and associates with a mesenchymal cell-state (9). Similarly, the receptor tyrosine kinase AXL can confer resistance to both osimertinib and erlotinib (10,11). It has also been associated with EMT in NSCLC and its concurrent inhibition with osimertinib can prevent the growth and delay the relapse in NSCLC patient-derived xenografts (PDXs) (5,9,12). Clinical case reports have also proposed RB1 loss and lineage switch to enable acquired resistance to anti-PD-1 antibody treatment (13,14). Consistent observations have been made in the first-line treatment of EGFR-mutant NSCLC with osimertinib, where PD-L1 expression has been shown to be a negative prognostic factor, and PD-L1 expression > 50% was associated with up to two-fold risk of death or progression, suggesting the role of PD-L1 role in osimertinib resistance (13,15–17). However, the underlying mechanisms driving these clinical observations have not yet been identified. Therefore, it remains to be elucidated whether these associations among different axes of plasticity – EMT, lineage switch from LUAD to LUSC, osimertinib resistance, PD-L1 upregulation, and reduced RB1 activity – are merely co-co-occurring or correlative processes, or whether they can causally drive and reinforce one another.

Here, we integrate dynamical simulations of an underlying regulatory network with extensive analysis of bulk, single-cell and spatial transcriptomic data of osimertinib-treated or EMT-induced cells, to demonstrate that EMT activation, PD-L1 upregulation, reduced RB1 activity interconnected with osimertinib resistance can reinforce each other due to the presence of higher-order feedback loops in the network, forming two mutually inhibiting ‘teams’ of nodes. Perturbing one axis can drive coordinated changes along many other axes too. This coordinated expression of EMT, PD-L1, AXL and reduced RB1 activity – enabled by the ‘teams’ topology – is also observed in patient samples. Finally, we report that such coordinated activation of more than one axes – EMT, upregulation of AXL or PD-L1 and defective RB1 pathway activity – lead to worse clinical outcomes as compared to their individual roles, underscoring that such reinforcement can amplify the fitness of NSCLC cells under therapeutic stress. Our results suggest breaking these ‘teams’ of nodes as a network topology-driven potential therapeutic strategy.

## Results

### Gene regulatory network depicting a ‘teams’ structure and an association between osimertinib resistance and EMT

We first identified a minimal gene regulatory network (GRN) that integrates the known interactions among key molecular players of EMT (ZEB1, SNAIL, miR-200, miR-34), AXL, RB1 and PD-L1 (**Fig 1A**). We first identified a minimal gene regulatory network (GRN) that integrates the known interactions among key molecular players of EMT (ZEB1, SNAIL, miR-200, miR-34), AXL, RB1 and PD-L1 (Fig 1A). AXL activation can enhance the survival of DTPs and its inhibition during either the initial or tolerant phases can delay tumour re-growth compared to osimertinib alone (10). Clinically, its high expression associates with a low response rate to EGFR-TKI, and GAS6-AXL signalling is upregulated in EGFR-treated patients presenting with residual disease (10,18). AXL can activate NF-κB signalling that can stabilize SNAIL (19), thus driving EMT. AXL also activates miR-34a that can repress EMT (20), while miR-34a can inhibit AXL, forming a negative feedback loop (18). SNAIL and miR-34 form a mutually inhibitory feedback loop (21), similar to that reported between ZEB1 and miR-200 family (22). RB1 can repress ZEB1 (23), thus RB1 loss can lead to induction of ZEB1 (24). Conversely, ZEB1 can inactivate RB1, forming yet another positive reciprocal loop (25). While wild-type p53 can inhibit NF-κB (26), NF-κB activates p53 in response to stress (27), constituting a negative feedback loop. Phosphorylated RB can suppress NF-κB transcriptional activity and its downstream targets such as PD-L1 (28); inactivation of RB can promote pro-inflammatory signals in many cancers (29). NF-κB can activate miR-192-5p (30) whose over-expression can decrease RB1 mRNA and protein levels in lung cancer cells (20). These intertwined positive and negative feedback loops among different molecules obviate any intuitive understanding of the phenotypic landscape enabled by this GRN. Thus, we developed a mechanistic dynamical model capturing the above-mentioned interactions to elucidate the emergent dynamics of this GRN.

**Figure 1.**
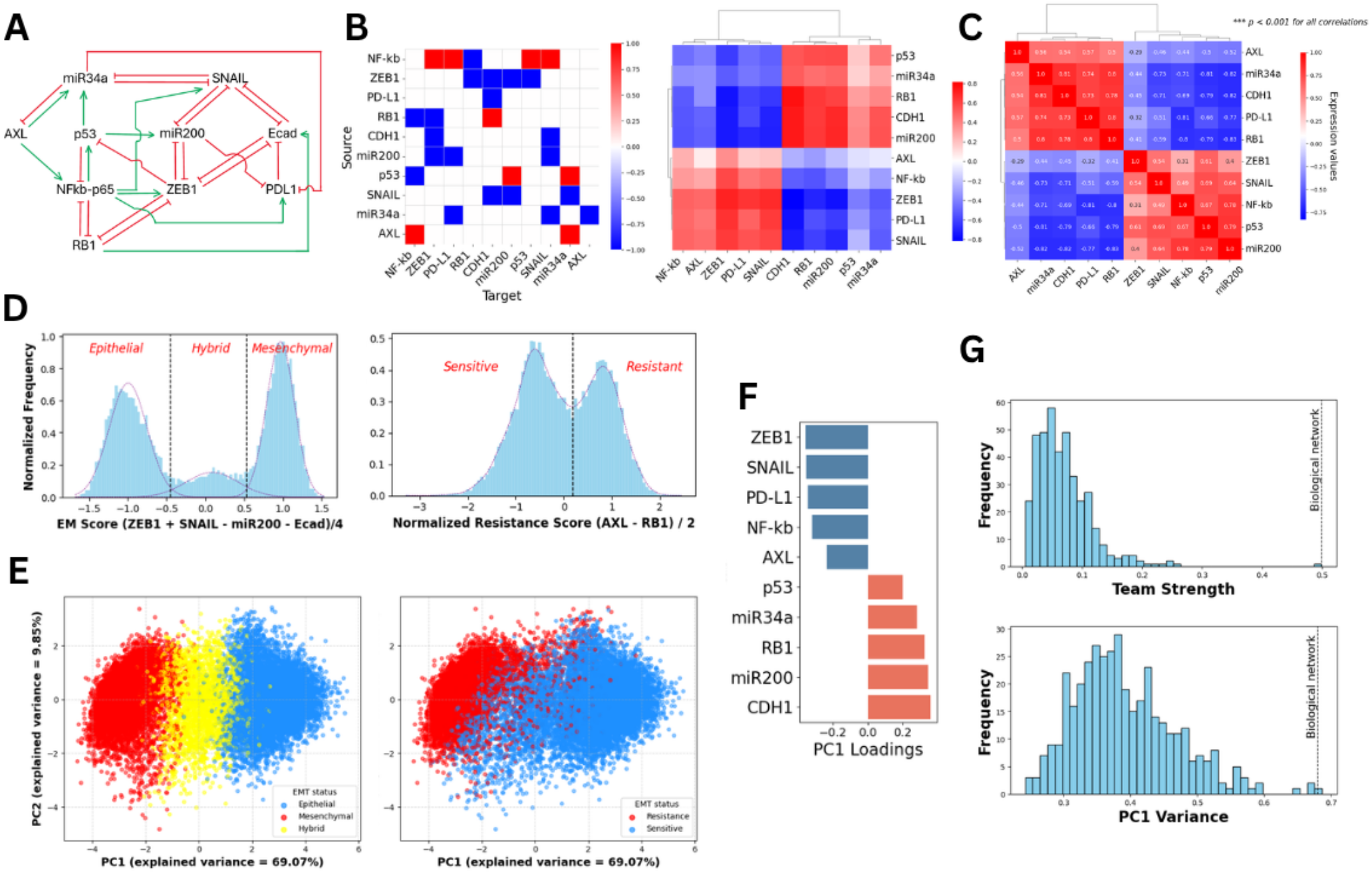
Dynamical simulations of EMT-RB1-AXL gene regulatory network (GRN). (A) GRN showing signaling among EMT markers, RB1 and AXL; Green colored edges represent activation, and red colored edges represent inhibition. (B) Adjacency matrix representing +1 as activation links and -1 as inhibition links with influence matrix (l=10) depicting “teams” in the network topology. (C) Correlation matrix of all possible steady states from RACIPE simulations of the network showing similar teams as shown in influence matrix. (D) Normalized frequency density histogram of EMT score (= (mir200 + Ecad – ZEB1 – SNAIL)/4) fitted to Gaussian mixture models and Resistance score (= (AXL – RB1)/2)) fitted to Kernel density estimation. (E) Principal Component Analysis (PCA) dimensionality reduction plots of all possible steady states color coded by phenotypes derived from density histograms. (F) PC1 loadings corresponding to each node in the network showing same signs for members in each team. (G) PC1 variance and team strength frequency histogram showing biological network with highest team strength and PC1 variance.

These interactions can be represented in the form of an adjacency matrix (31), where each row depicts the source, each column denotes the target, and each cell can take 3 values: -1 (red) indicating an inhibition edge, +1 (blue) indicating an activation edge, and 0 (white) for no direct regulation (**Fig 1B, left**). The adjacency matrix can be then used to generate an influence matrix, where each cell denotes the net impact that one node (source) in the GRN has on another one (target), through multiple paths of varying lengths containing multiple activation and/or inhibition edges in each path (**Fig 1B, right, S1A**). Thus, the influence matrix captures long-range regulatory interactions in a GRN (31). The influence matrix for this GRN revealed the presence of two “teams” such that members within each team effectively activate one another, while those across the two teams effectively inhibit each other to varied extents. One ‘team’ comprises E-cadherin, p53, miR-200, miR-34 and RB1 – all of which are known to maintain or drive an epithelial phenotype (32–36). The other ‘team’ consists of ZEB1, PD-L1, SNAIL, AXL and NF-κB – all of which can drive a mesenchymal immunosuppressive cell-state (32,37–40). Next, we computed the team strength that is defined on a scale of 0 to 1 and quantifies the degree of separation between the two teams. The team strength for this influence matrix was found to be 0.49. Collectively, these observations suggested that this GRN topology is structured to enable the presence of two mutually inhibiting and self-reinforcing ‘teams’ of nodes that can drive multiple axes of cellular plasticity.

Further, we simulated the dynamics of this GRN using RACIPE – a computational tool that simulates the emergent dynamics of a coupled set of ODEs representing a GRN over multiple initial conditions and biologically relevant kinetic parameter sets – to identify its different possible steady-states (41). We obtained over 25,000 unique steady state values across 10,000 parameter sets, indicating the prevalence of multi-stability (co-existence of different steady-states/phenotypes) as a feature of this GRN. Correlation heatmap of this ensemble of phenotypes shows the same ‘team’ structure as observed in the network topology, with miR-200 and RB1 on one team, and ZEB1 and AXL on the other team (**Fig 1C**). To decode the functional relevance for the steady-states, we calculated corresponding EMT scores ( (= ZEB1 + SNAIL – miR-34 – miR-200)/4 ) and resistance scores ( = (AXL – RB1)/2 ), given the antagonistic roles of EMT-inducing factors (ZEB1 and SNAIL) and EMT-inhibiting microRNAs (miR-200 and miR-34) in regulating EMT (21,42), and those of RB1 and AXL in driving osimertinib resistance (9,10). A normalized frequency histogram of EMT scores revealed trimodality, with two dominant peaks corresponding to epithelial and mesenchymal phenotypes. Similarly, the histogram of resistance scores showed two peaks corresponding to sensitive and resistance phenotypes (**Fig 1D**). To better visualise the various phenotypes enabled by this GRN, we projected the steady-states using principal component analysis (PCA) and observed that the epithelial cluster was more likely to be sensitive, and mesenchymal cluster was much more likely to be resistant. Conversely, the sensitive cluster was more likely to be epithelial, and resistant one was more likely to be mesenchymal (**Fig 1E, S1C**). The loading coefficients of principal component 1 (PC1) further confirmed this association that all players inhibiting EMT or osimertinib resistance had the same sign, while those driving EMT or osimertinib resistance had the opposite sign (**Fig 1F**), further validating the composition of ‘teams’ of nodes observed in the influence matrix and pairwise correlation matrix for this GRN.

To examine whether the emergence of these teams is specific to this GRN topology, we generated 400 randomized network topologies while preserving the number of total activating and inhibitory edges in the network but randomly shuffling individual regulatory edges. We performed RACIPE simulations on each randomized network. A scatter plot comparing the variance of the first principal component (PC1) of steady-state expression profiles with the corresponding team strength across network topologies revealed that this GRN had both the highest PC1 variance and the strongest team strength (**Fig 1G, S1D**). This observation suggests that the topology of the underlying biological network is evolutionarily tuned to coordinate phenotypic plasticity across multiple behavioral axes through by maintaining the “teams” structure.

To model the impact of frequent genetic mutations on this phenotypic landscape, we modified the GRN based on experimentally reported links related to p53 gain-of-function (GOF) mutations such as R175H, R273H, R280K and R248W. R175H-mutant p53 can constitutively activate NF-κB signaling, leading to more aggressive behaviors (43), and disrupt activation of miR-200 and miR-34a, thus promoting EMT and stem cell properties (44). This modified network had similar emergent dynamics with trimodal EMT scores and bimodal resistance scores but with mutant-p53 (p53-m) changing teams from being in the epithelial one to being in a mesenchymal one (**Fig S2D**). We noticed an increased proportion of hybrid epithelial/mesenchymal (E/M) phenotypes in EMT score distribution (**Fig S2C**).

Overall, the GRN we constructed based on individual experimental data about interconnections among the nodes in the GRN revealed – based on topological and dynamical attributes – that EMT and osimertinib resistance in NSCLC operate in an interconnected and coordinated manner. The interconnections manifest as two functional “teams” - epithelial team (miR-200, miR-34a, CDH1, RB1) linked to osimertinib-sensitivity, while the mesenchymal team (ZEB1, SNAIL, AXL) with osimertinib-resistance.

### Clinical bulk transcriptomic data is consistent with the composition of these ‘teams’

To assess whether the predicted association between EMT and osimertinib resistance obtained by GRN modeling holds true in NSCLC cell lines and patient tumors, we analyzed bulk transcriptomics data across multiple contexts. We started with NSCLC cell lines in the Cancer Cell Line Encyclopedia (CCLE) (45) and performed single-sample Gene Set Enrichment Analysis (ssGSEA) (46) for relevant gene signatures – a) Hallmark EMT from the Molecular Signature Database (MSigDB) (47), b) separate epithelial and mesenchymal signatures for cancer cell lines (KS-Epi, KS-Mes) (48), and c) pan-cancer PD-L1 associated signature from our prior study, based on genes most strongly correlated with PD-L1 expression levels across 13 cancer types (49). Previously reported gene signatures were used to characterize RB1-associated transcriptional programs: a) biallelic RB1 loss signature – a pan-cancer signature based on 995 CCLE cell lines (50), b) defective RB1 signature predicting deleterious RB1 mutations or dysregulation of the RB pathway, derived from pan-cancer proteogenomic datasets (51), and c) RB1 pathway activity signature representing the downstream transcriptional output of CDK4/6-RB axis across 31 TCGA cancer types (52), and d) RB1 loss signature of genes upregulated by RB1 deletion in murine liver or in fibroblastic models (53).

Across NSCLC cell lines, as expected based on the reported E-M spectrum in NSCLC (54,55), we observed the KS-Epi and KS-Mes ssGSEA scores to be negatively correlated (r = - 0.77, p < 0.001) (**Fig 2A, left**). AXL expression was positively correlated with ssGSEA scores for Hallmark EMT signature (r = +0.79, p < 0.001) (**Fig 2A, middle**) and with those for PD-L1 associated signature and NF-κB pathway (r = +0.43, p < 0.001) (**Fig S3A**). ssGSEA score for genes downregulated in defective RB1 signature was positively correlated with KS-Epi ssGSEA scores (r = +0.58, p < 0.001) (**Fig 2A, right**), but negatively with EMT as measured by Hallmark EMT or KS-Mes ssGSEA scores (Fig S3A). Conversely, ssGSEA scores for genes upregulated in defective RB1 signature correlated positively with those overexpressed during biallelic RB1 loss but negatively correlated with RB1 pathway activity and with KS-Epi ssGSEA scores (**Fig S3A**). All these trends endorse the ‘teams’ composition we observed in the GRN – EMT being associated with reduced RB1 activity and increased AXL levels.

**Figure 2.**
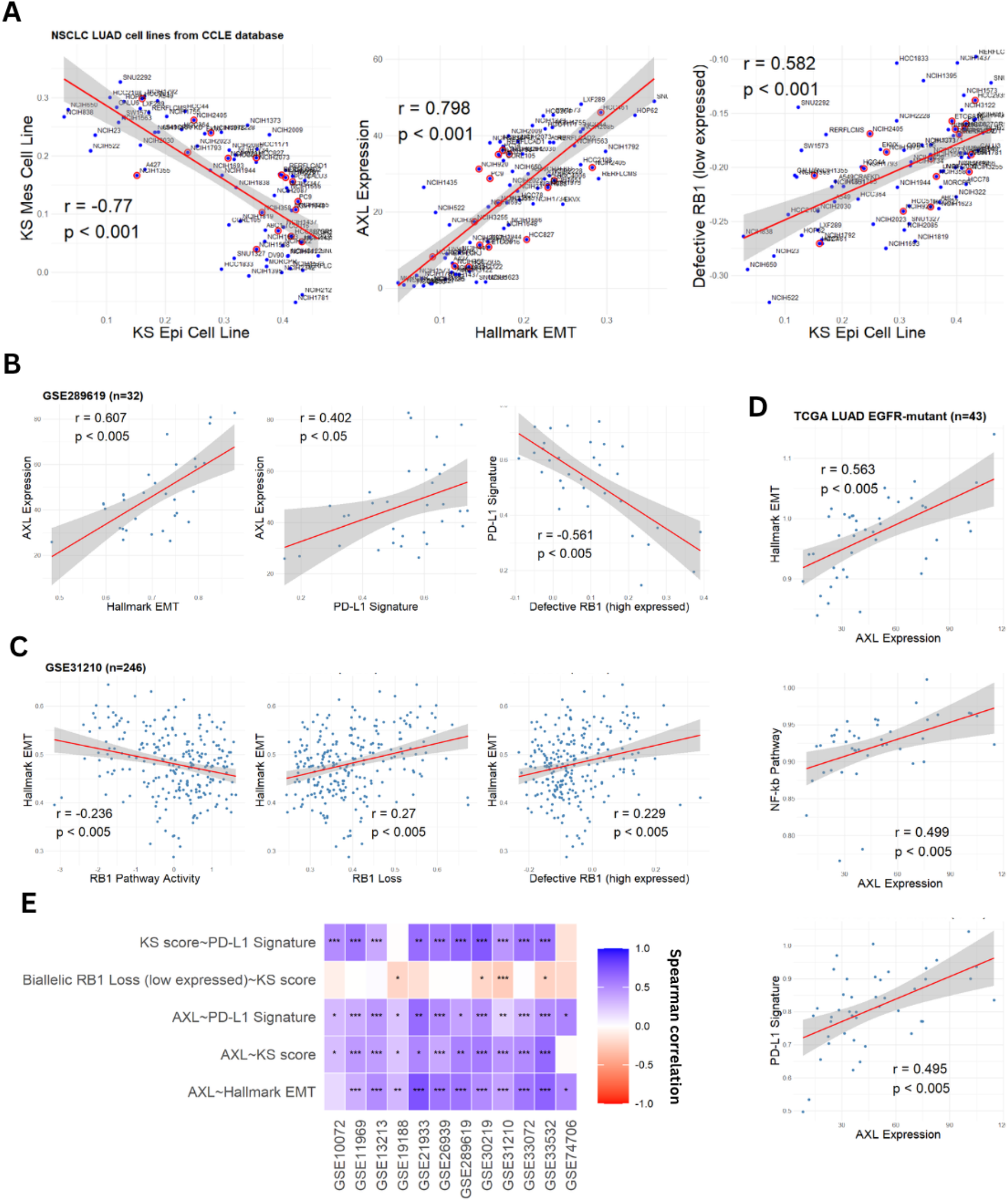
Bulk transcriptomic analysis for lung cancer patient and cell line datasets. (A) Correlation of activity of Epithelial-Mesenchymal signatures, Hallmark EMT activity – AXL gene expression and Hallmark EMT – RB1 pathway activity (ssGSEA scores) across all lung cancer cell lines depicting the baseline behaviour of these cell lines. (B) Scatter plots of ssGSEA scores of EMT-associated signatures, RB1-associated signatures and AXL gene expression for 32 patients with Osimertinib (OSM) monotherapy. (C) Scatter plots for LUAD patients with predominant EGFR mutant population, GSE31210 (D) Scatter plots of AXL expression with EMT, PD-L1 and NF-κB signatures for TCGA-LUAD patients with EGFR mutations (E) Heatmap of correlations between different signature activities and gene expressions for patient bulk transcriptomic datasets.

We next investigated RNA-seq data for 32 EGFR-mutant NSCLC patients treated with osimertinib as the first-line therapy (GSE289619) (56). AXL expression was found to correlate positively with ssGSEA scores for PD-L1 associated signature (r = +0.40, p < 0.05) and for Hallmark EMT signature (r = +0.60, p < 0.001) (**Fig 2B**). PD-L1 signature was negatively correlated with genes upregulated in defective RB1 signature (r = -0.56, p < 0.001) (**Fig 2B**). In a cohort of stage I and II LUAD patients (GSE31210) (57), Hallmark EMT scores correlated positively with RB1 loss signatures (r = +0.27, p <0.05) and with genes upregulated in defective RB1 cases (r = +0.23, p < 0.05), but negatively with RB1 pathway activity (r = -0.23, p < 0.005) (Fig 2C). In TCGA LUAD EGFR-mutant patients (n=43), AXL expression was positively correlated with ssGSEA scores for PD-L1 associated (r = +0.495, p < 0.001), NF-κB signaling (r = 0.49, p < 0.001) and Hallmark EMT signatures (r = +0.563, p < 0.001) (Fig 2D). In both these cohorts, EMT correlated positively with AXL expressions and with ssGSEA scores for PD-L1 associated and NF-κB signaling signatures (Fig S3B-C), endorsing a consistent relationship between EMT and osimertinib resistance, as predicted by GRN simulations.

We next investigated additional LUAD or NSCLC patient cohorts – GSE10072 (58), GSE11969 (59), GSE13213 (60), GSE19188 (61), GSE21933 (62), GSE26939 (63), GSE30219 (64), GSE33072 (65), GSE33532 and GSE74706 (66) –where AXL expression was consistently positively correlated with ssGSEA scores for PD-L1 associated signatures, NF-κB signaling, Hallmark EMT genes (**Fig 2E**).

Collectively, these results across multiple cell lines and patient cohorts show the pairwise correlations between AXL overexpression, low or loss of RB1 activity, and mesenchymal transcriptional programs with elevated PD-L1– associated signatures. These NSCLC cell line and EGFR-mutated patient cohort data support the prediction from our GRN simulations on the composition of two ‘teams’ – mesenchymal/resistant team marked by EMT, AXL overexpression and immune signaling (PD-L1/NF-κB) vs. epithelial/sensitive team marked by intact RB1.

### Perturbation experiments reveal bidirectional regulation of EMT and osimertinib resistance

Correlations observed in above-mentioned datasets do not necessarily imply causation. To assess whether the perturbation of one phenotypic axis influenced the behavior along other axes, we focused on datasets connected to perturbation experiments where NSCLC EGFR-mutant cells were exposed to growth factors such as TGF-β to induce EMT or treated with osimertinib.

First, in HCC827 cells (characterized by EGFR exon 19 deletion) induced to undergo EMT via TGF-β treatment (GSE49644) (67), the genes highly expressed in biallelic RB1 loss or defective RB1 signature were upregulated (**Fig 3A, i**), indicating an osimertinib-resistant phenotype. Similar results were observed in HCC827 cells with ZEB overexpression (GSE81167) (68) suggesting a core regulatory network at play connecting EMT to EGFR-TKI resistance (**Fig 3A, ii**). These trends were not observed in TGF-β treated A549 cells which do not harbor any EGFR mutations (**Fig S4**). Conversely, in experiments involving 21-day treatment with osimertinib to investigate drug-tolerant persisters (DTPs) in HCC827 and HCC2935 (characterized by EGFR exon 19 deletion) cells (GSE193258) (65), we observed higher AXL expression and upregulated EMT, and downregulation of genes that have low expression in the biallelic RB1 loss case (**Fig 3B**). DTPs are a small subpopulation of cancer cells that survive treatment with standard-of-care therapies without acquiring stable genetic resistance, but via reversible non-genetic mechanisms (69). A short (24 hours) or long (4 or more days) washout corresponding to drug withdrawal, until the exponential cell proliferation resumed, however, did not lead to a decrease in AXL expression or EMT ssGSEA scores, suggesting putative mechanisms of cellular memory, as observed experimentally for EMT (70,71).

**Figure 3.**
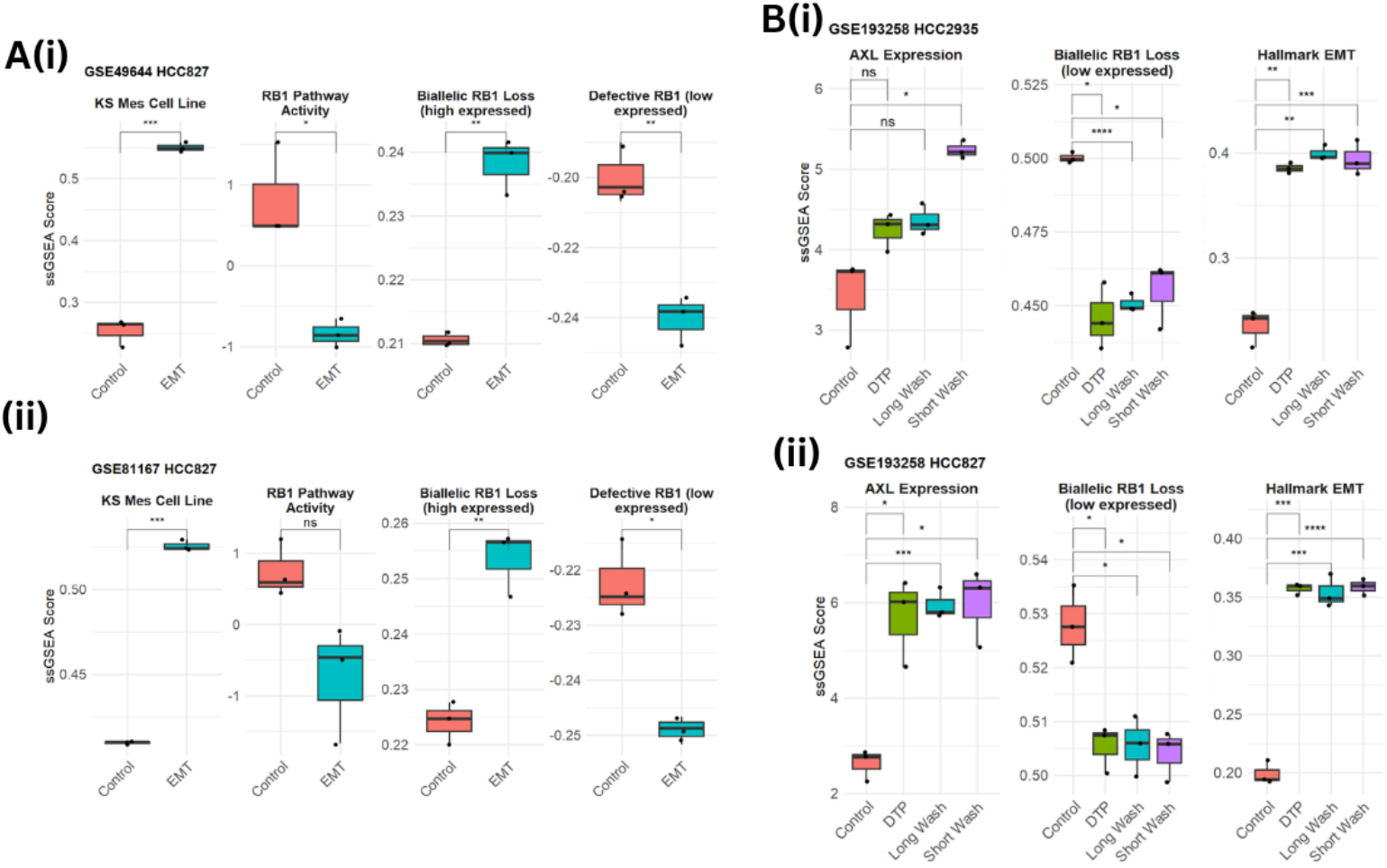
Gene expression analyses in perturbation experiments for different cell lines. (A(i)) Box plot of enrichment scores in TGF-β EMT induction experiment in HCC827 cell lines depicting EMT induction leading to upregulation of RB1 loss programs, GSE49644; (ii) Same as (i) but EMT induction performed by ZEB1 overexpression in HCC827 cell line, GSE81167. (B) Box plot of enrichment scores for Hallmark EMT, RB1 associated signatures, and AXL expression levels upon OSM exposure followed by drug holiday (short and long wash groups) in HCC2935(i) and HCC827(ii), GSE193258, depicting chronic EMT induction in DTPs.

Overall, the analysis of these *in vitro* perturbation datasets reveals that EMT and osimertinib resistance are not only correlated but are mechanistically coupled through reciprocal feedback loops in the underlying GRN. This bidirectional regulation underlies the consistent correlation pattern observed in patient cohorts, enabling stable and self-reinforcing resistant states.

### RB1 loss, AXL and EMT programs are correlated at a single-cell and spatial transcriptomic level

Following the pairwise correlation and perturbation analysis at bulk transcriptomic level, we moved to single-cell and spatial transcriptomic data from patients to further examine the associations between RB1 loss, AXL expression and activation of EMT.

We analyzed single-cell RNA-seq (scRNA-seq) and spatial transcriptomics (ST) from patient samples belonging to different stages of LUAD – from adenocarcinoma in situ (AIS) to minimally invasive adenocarcinoma (MIA) and invasive adenocarcinoma (IAC) (GSE189357) (72). In scRNA-seq data, the activation of EMT was negatively correlated with RB1 activity signatures (r = -0.39, p < 0.001 for AIS; r = -0.80, p < 0.001 for MIA; r = -0.69, p < 0.001 for IAC) (**Fig4A**), and positively correlated with AXL expression levels (r = +0.41, p< 0.001 for AIS; r = +0.55, p < 0.001 for MIA; r = +0.66, p < 0.001 for IAC) (**Fig S5A**) across different stages. AXL expression was moderately negatively correlated with RB1 pathway activity (r = -0.3, p < 0.001) in AIS, and the trend was stronger in IAC (r = -0.49, p < 0.001), following the patterns for EMT activation (**Fig S5A**). In AIS and MIA stages, the PD-L1 associated signature was strongly positively correlated with EMT (r = +0.42, p < 0.001 for AIS; r= 0.52, p < 0.001 for MIA), NF-κB activity (r = +0.6, p< 0.001 for AIS; r = +0.48, p< 0.001 for MIA) and with the genes down-regulated in defective RB1 case (r = +0.15, p< 0.001 for AIS; r = +0.45, p< 0.001 for MIA) (**Fig S5A**). These trends highlight that EMT, AXL, PD-L1 activity, and RB1 loss signatures can be correlated at a single-cell level.

For spatial transcriptomic (ST) data from these patients (GSE189487), we performed GSEA across spatial spots using AUCell (73). Starting with patient 1 belonging to IAC, we observed a clear similarity in the spatial activity levels of Hallmark-EMT and KS-Mes gene-sets (r = +0.6, p < 0.001); both of them were positively correlated with AXL signature (r = +0.39, p < 0.001 for Hallmark EMT; r = +0.64, p < 0.001 for KS-Mes). The spatial activity pattern of KS-Epi gene set was antagonistic to EMT patterns (r = -0.16, p< 0.001 for Hallmark EMT; r= -0.44, p < 0.001 for KS-Mes) (Fig 4B, C). However, the spatial activity of PD-L1 associated signature was weakly correlated to EMT (r = +0.16, p< 0.001 for KS-Mes; r = +0.02; p > 0.05 for Hallmark EMT) and AXL (r = +0.22, p < 0.001) (Fig 4C). Moreover, while the RB1 activity pathway was positively correlated with KS-Epi (r = +0.31, p < 0.001), it was also positively correlated with EMT (r = +0.39, p< 0.001 for KS-Mes; r = +0.20; p > 0.05 for Hallmark EMT). One possible interpretation of this result can be the association of RB1 pathway with a partial EMT signature, reminiscent of trends seen for basal gene signature in breast cancer (74).

**Figure 4.**
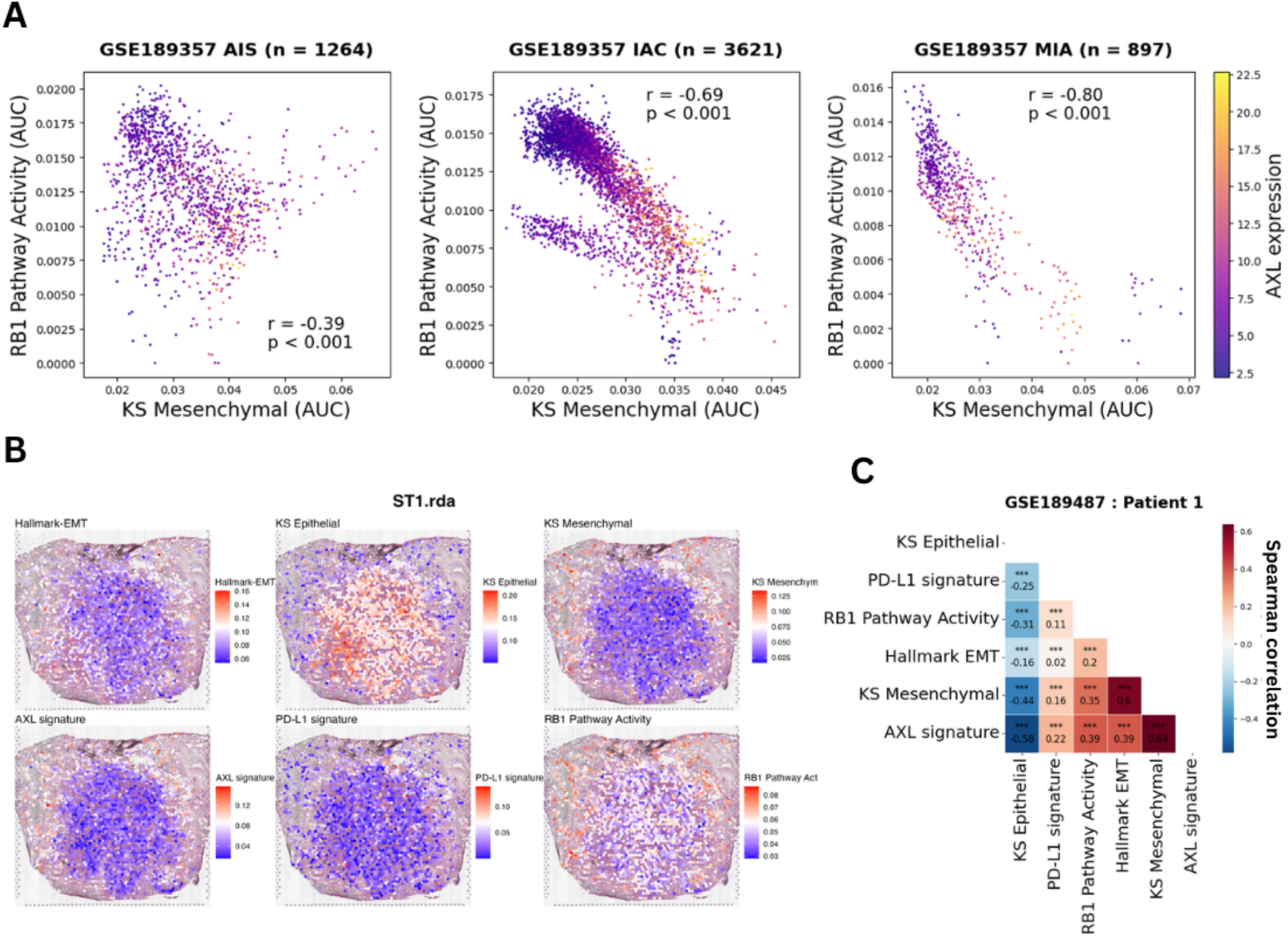
Single cell RNA-seq analysis for LUAD patient datasets from different clinical stages. (A) AUC scores correlation plots showing negative correlations between mesenchymal and RB1 pathway signatures colored by AXL expression values from less invasive (AIS) samples to minimally invasive (MIA) and highly invasive (IAC) samples (GSE189357). (B) Spatial transcriptomics analysis of Patient-1 corresponding to IAC (GSE189487) with gene set enrichment analysis showing strong AXL signature and EMT score correlations.

Further analyses of more patients from this data set – Patient 2 (IAC), Patient 6 (MIA) and Patient 8 (AIS) – suggests that correlation trends between Hallmark-EMT, KS-Mes, KS-Epi and AXL is conserved regardless of the clinical stage of the patient, while RB1 and PD-L1 signature activity and their association with other signatures studied may vary depending upon the clinical stage (**Fig S5B**). Analysis of ST of four tumor samples from another dataset (E-MTAB-13530 (75)) showed consistent association between different genesets as we noted previously. KS-Mes, Hallmark EMT and AXL are strongly positively correlated with one another and negatively correlated with KS-Epi geneset (**Fig S6**).

These observations reinforce our observations that the coordinated relationship between EMT, AXLS activation and RB1 loss is maintained at single-cell and spatial levels, supporting a conserved, multi-axis program of tumor plasticity and osimertinib resistance.

### Coordinated plasticity axes correlate with worse patient outcomes across multiple cohorts

We next asked whether this coordinated response across many axes of plasticity is clinically relevant. To address this, we performed Kaplan Meier analysis and Cox-hazard ratio analysis for various gene signatures taken alone or in combination, for three publicly available LUAD cohorts with survival metrics – GSE31210 (57), GSE68465 (76) and the TCGA-LUAD cohort. We segregated patients into two groups based on median value of each feature and then compared the survival between them.

In GSE31210, patients with lower RB1 pathway activity had worse overall survival (OS) as compared to those with higher activity (HR = 3.69, p < 0.01) (**Fig 5A**). Similarly, patients with high activity levels of the geneset highly expressed in defective RB1 cases had worse OS (HR = 3.29, p < 0.01) (**Fig 5B**). Consistent with our ‘teams’ composition and pairwise correlation trends, high expression of PD-L1 (CD274) (HR = 2.03, p < 0.05) or high activity of Hallmark EMT signature (HR = 2.13, p < 0.05) was associated with worse OS (**Fig 5D, E**). However, the trends were not significant for AXL expression or the signature of genes that are expressed at a lower level in defective RB1 cases (**Fig 5C, F**).

**Figure 5.**
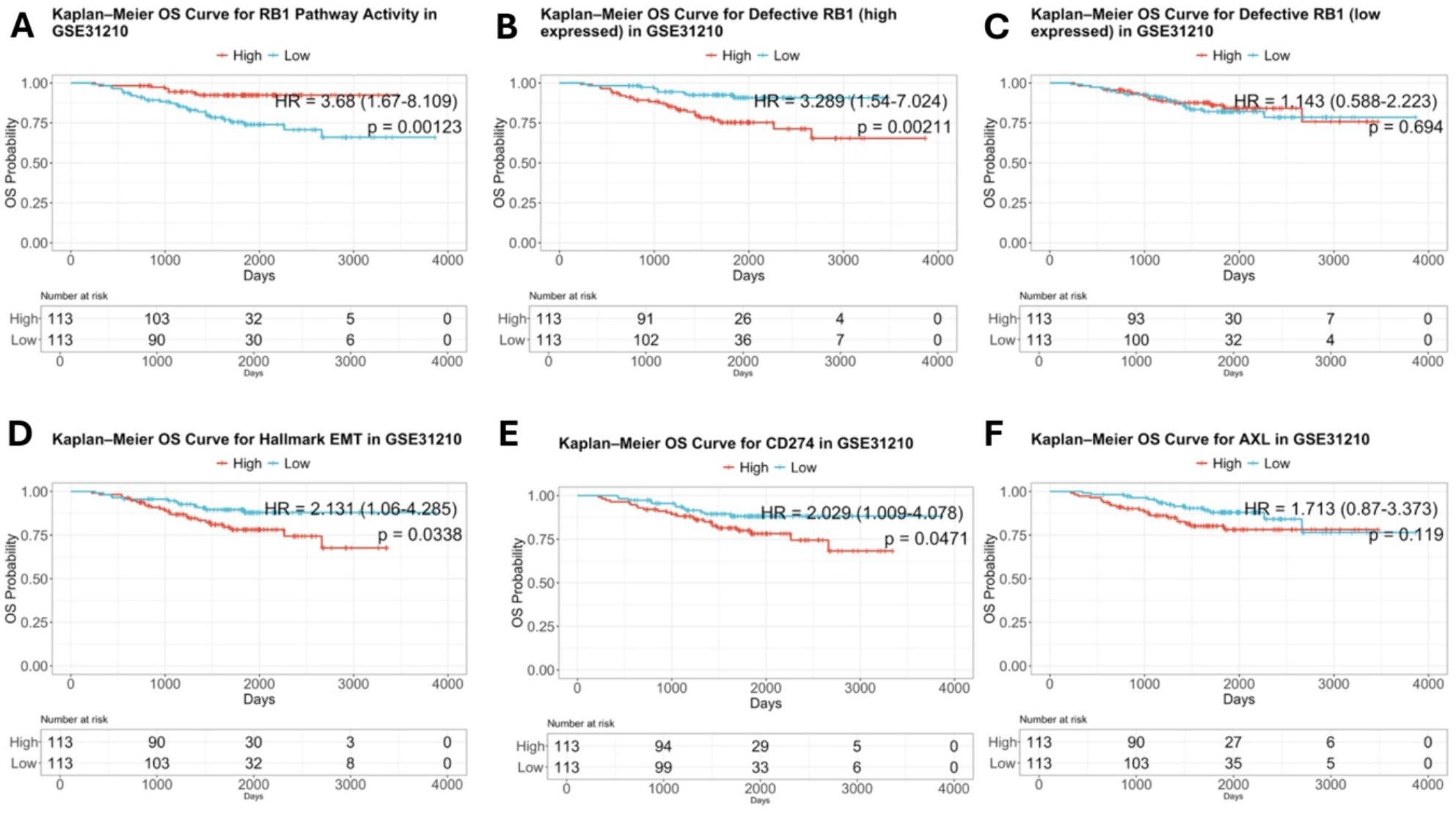
Overall Survival Kaplan-Meier (KM) plots for enrichment scores of different gene sets and expression levels of genes in GSE31210 patient cohort separated by median. (A) Overall Survival KM plots for activity levels of Hallmark-EMT; (B, C) Overall Survival KM plots for expression levels of CD274 (PD-L1 gene) and AXL genes; (D, E) Overall Survival KM plots for activity for levels of Defective-RB1 (high expressed) and Defective-RB1 (low expressed) gene sets. (OS: Overall Survival)

Beyond OS, we also calculated other trial endpoints - disease-free survival (DFS), defined as the time from start of treatment with curative intent to disease recurrence or death, disease-specific survival (DSS), defined as the time from treatment start to death specifically caused by the disease, and progression-free survival (PFS), defined as the time from treatment start with the intent of tumour shrinkage to disease progression or death from any cause. All the previous trends were recapitulated in the DFS metrics for GSE31210 dataset as well (**Fig S7**), as well as OS and DFS for GSE68465 (**Fig S8**), and for OS, DFS, DSS and PFS for the TCGA-LUAD cohort (**Fig S9**). For the TCGA-LUAD cohort, the low scores for the signature of defective RB1 (low expressed) genes also associated with worse patient outcomes, irrespective of the survival metric looked at – OS, DSS, DFS and PFS (**Fig S9C, i-iv**). Together, these trends indicate that activation of these individual axes of plasticity – defective RB1, EMT or PD-L1 upregulation – can be sufficient to drive worse outcomes.

Motivated by these findings, we interrogated whether simultaneous activation of these plasticity axes – as enabled by the ‘teams’ of nodes – can further aggravate the survival probability as compared to individual plasticity axes being activated. We probed various such combinations – taken two at a time – of a) activity levels of Hallmark EMT signature, b) AXL expression levels, c) expression levels of PD-L1 gene (*CD274*), d) activity levels of RB1 pathway, e) activity levels of the defective RB1 (high expressed) geneset, and f) activity levels of defective RB1 (low expressed) geneset. Given the well-established association among Hallmark EMT signatures, expression levels of AXL and CD274, and the *RB1* loss signature, we took for reference the patient samples with low activity of Hallmark EMT and of defective RB1 (high expressed) signatures, low expression levels of AXL and PD-L1, and with high activity of RB1 pathway and of defective RB1 (low expressed) signatures.

For the heatmaps denoting OS metrics for different combinations of Hallmark EMT and RB1 pathway activity, we noticed that reduced activity of RB1 pathway alone could lead to worse OS (HR = 3.52, p < 0.05). Importantly, reduced activity of RB1 pathway along with increased activity of Hallmark EMT led to even worse outcomes (HR = 5.55, p < 0.01) (**Fig 6A, i**). Similarly, simultaneous activation of high Hallmark EMT and defective RB1 (high expressed) signatures led to worse outcomes as compared to either or none of these two signatures being upregulated (**Fig 6A, ii**), underscoring the synergistic role of the simultaneous activation of these two axes of plasticity in enabling aggressive clinical outcomes. Further, in another combination of AXL expression and RB1 pathway activity, we noticed that reduced activity of RB1 pathway led to worse outcomes (HR = 4.52, p < 0.05), but the worst survival was again shown in patient samples with reduced RB1 pathway activity and with increased AXL expression (HR = 6.57, p < 0.01) (**Fig 6B, i**). Reinforcing trends were recapitulated in other pairwise combinations too: a) AXL expression with defective RB1 (high expressed) signature (HR = 6.58, p < 0.05) (**Fig 6B, ii**), c) CD274 expression with RB1 pathway activity (HR = 6.70, p < 0.01) (**Fig 6C, i**), d) CD274 expression with defective RB1 (high expressed) signature (HR = 6.60, p < 0.01) (**Fig 6C, ii**), e) AXL expression with Hallmark EMT signature (HR = 2.71, p < 0.05), f) CD274 expression with Hallmark EMT signature (HR = 3.23, p < 0.05), g) AXL expression with CD274 expression (HR = 2.70, p < 0.05) (**Fig 6D**), h) PD-L1 signature with RB1 pathway (HR = 5.76, p < 0.01) and i) PD-L1 signature with defective RB1 (high expressed) (HR = 5.68, p < 0.01) (**Fig S11**). These abovementioned trends, along with those for pairwise combinations of defective RB1 (low expressed) with Hallmark EMT, AXL expression or CD274 expression, were distinctly observed for DFS too in this dataset (**Fig S10**). These co-expressed axes of phenotypic plasticity corresponding to the composition of ‘teams’ in the GRN – increased activity of EMT pathway, higher expression of AXL or CD274, and higher activity of defective RB1 (high expressed) or lower activity of defective RB1 (low expressed), reduced RB1 pathway activity – associated with worse OS and DFS in GSE31210 (**Fig S10**) as well as with worse OS, DFS, DSS and PFS in the TCGA – LUAD dataset (**Fig S13**), as compared to only one axis of plasticity being activated.

**Figure 6.**
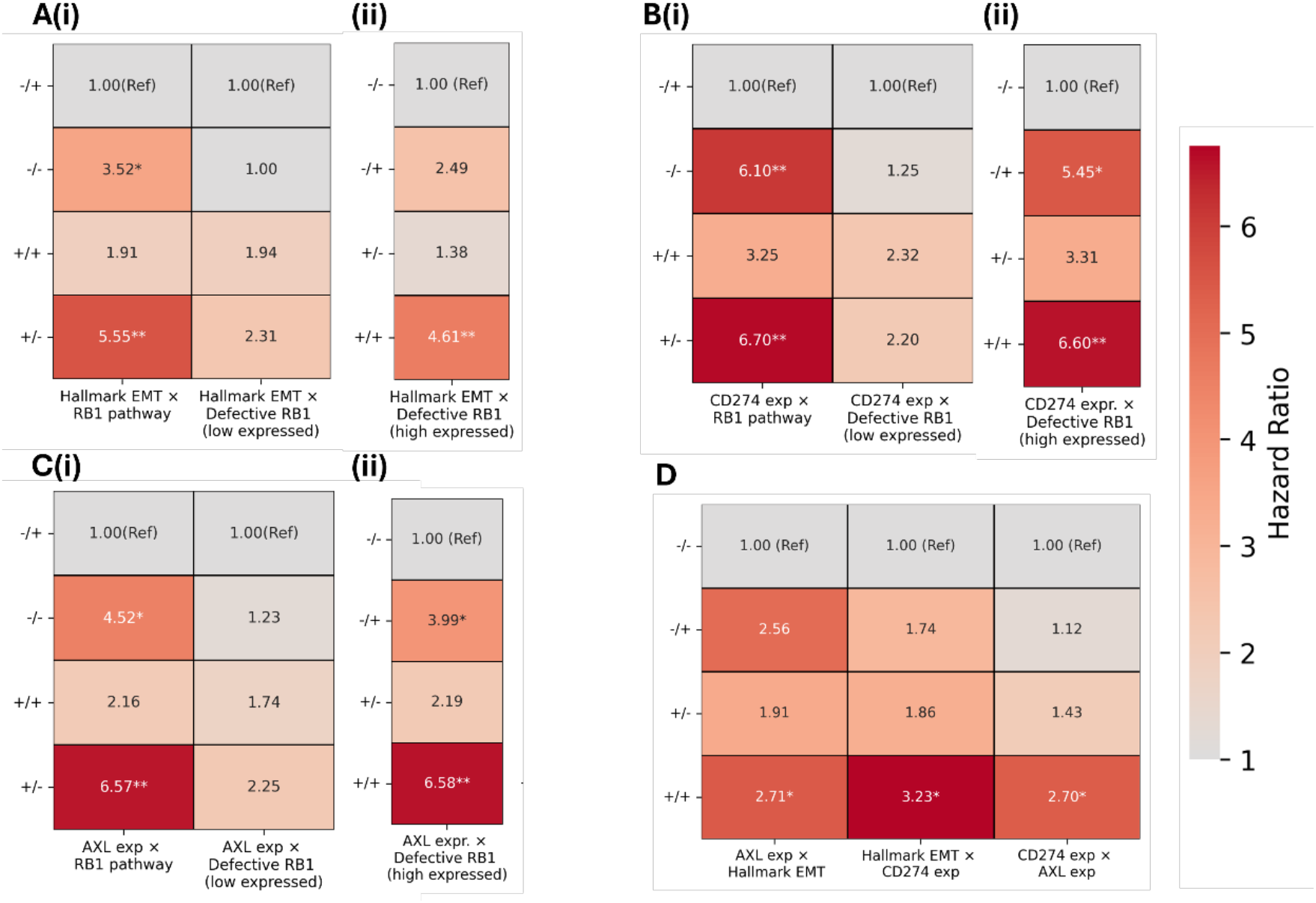
Overall Survival heatmaps showing Hazard Ratio (HR) values for double combinations of enrichment scores of different gene sets and expression levels of genes in GSE31210. (A) HR values for different combinations of Hallmark EMT vs RB1 Pathway vs Defective-RB1-loss (low expressed) and (ii) Defective-RB1-loss (high expressed); (B) Same as (A) for different combinations of CD274 (PD-L1 gene) vs (i) RB1 pathway, Defective-RB1-loss (low expressed) and (ii) Defective-RB1-loss (high expressed); (C) Same as (A) for different combinations of AXL gene vs (i) RB1 Pathway, Defective-RB1-loss (low expressed) and (ii) Defective-RB1-loss (high expressed); (D) HR values for different combinations of AXL gene vs Hallmark-EMT, Hallmark-EMT vs CD274 (PD-L1 gene) and CD274 (PD-L1 gene) vs AXL gene. (* indicates p-value < 0.05, ** indicates p-value <0.01, *** indicates p-value <0.001; OS: Overall Survival)

Finally, we performed pairwise Cox-hazard ratio analyses for the combinations of three modules – a) Hallmark EMT, defective RB1 (high expressed) and CD274 expression, b) Hallmark EMT, defective RB1 (high expressed) and AXL expression, and c) Hallmark EMT, CD274 expression and AXL expression – in GSE31210. Again, simultaneous upregulation of all three pathways/gene expression levels led to much worse OS and DFS periods as compared to only one or two of them being upregulated together, with the HR values ranging up to 9.45 (95% CI = 2.83-31.66) for DFS, and up to 8.11 (95% CI = 1.83-35.98) for OS (**Fig S15**).

Collectively, these results emphasize the clinical relevance of simultaneous activation of multiple axes of phenotypic plasticity in terms of potentially amplifying the fitness of a subpopulation of cancer cells that can aggravate disease progression and drastically shorten OS and DFS duration for patients.

### Basal-shift markers associate with EMT and osimertinib resistance

Besides EMT, a partial shift from LUAD to LUSC has also been associated with osimertinib resistance (77). To assess any causative connections between EMT and this basal-shift, we extended the current version of our GRN to incorporate key lineage-specific regulators including NKX2-1, FOXA2, SOX2 and ΔNp63. FOXA1/2 and NKX2-1 (TTF-1) are canonical pulmonary lineage markers enabling lineage fidelity (78,79). NKX2-1 regulates the localization and activity of FOXA1/2, contributing to epigenetic reprogramming of cells (80). Functionally, SOX2 supports proliferation and preserves basal/stem-like identity, while NKX2-1 is essential for specifying alveolar epithelial fate (81,82). SOX2 and NKX2-1 can mutually repress each through promoter site binding (83). ΔNp63, a non-transactivating isoform of the p63 gene, also serves as a well-established marker of squamous/basal identity (84). LUAD marker NKX2-1 repress ΔNp63 through binding to distal enhancers in lung organoid culture (85), while SOX2 overexpression has been shown to mediate LUSC phenotype through p63 (both TP63 and ΔNp63 isoforms) upregulation (81).

We further looked at interactions of these lineage markers with the EMT markers, AXL and RB1 in the previous network. SOX2 downregulation has been implicated in inhibition of EMT, while FOXA2 suppresses EMT in NSCLC (86,87). NKX2-1 has also been studied to restrict metastasis through inhibition of mesenchymal markers SNAIL and SLUG during lung development and carcinogenesis (88). AXL downregulation has been shown to decrease SOX2 levels, while RB1 loss has been associated with increase in SOX2 levels in NSCLC (89,90). To restore the dilution of resistant-sensitive phenotypic distributions caused by earlier lineage marker interactions, we incorporated additional nodes representing apoptotic sensitizers in NSCLC cells, such as FOXO3a. Suppression of FOXO3a by NF-κB has been studied to contribute to gefitinib resistance, while FOXO3a itself can suppress AXL, SNAIL and SLUG via suppression of HIF-1α (91–93).

Dynamical simulations of this updated GRN reveal a bimodal distribution of LUAD-LUSC score (**Fig 7A, i-ii**) (where higher scores denote a LUSC-like state, and lower scores denote a LUAD-like state), while preserving a multi-modal distribution for the EMT scores and osimertinib resistance scores (**Fig S16A**). Conditional probability analysis suggests that the LUAD-like phenotype is more likely to be epithelial whereas LUSC-like state is more likely to be mesenchymal (**Fig 7A, iii**), and vice versa. Similarly, LUAD-like phenotype is more likely to be osimertinib-sensitive, while LUSC-like state is more likely to be osimertinib-resistant, and vice versa (**Fig 7A, iii**). The association between EMT and osimertinib resistance was maintained as for the earlier GRN (**Fig S16B**), thereby implying that the LUSC-like phenotype associates with osimertinib resistance based on emergent dynamics of this underlying GRN. Moreover, PC1 loadings from PCA analysis of steady states from this modified network results in Epi-sensitive-LUAD markers (p53, miR-34a, FOXO3a, CDH1, miR-200, RB1, NKX2-1, FOXA2) and Mes-resistant-LUSC markers (ZEB1, SNAIL, PD-L1, SLUG, AXL, HIF-1α, SOX2, ΔNp63) having same sign (**Fig 16C**).

**Figure 7.**
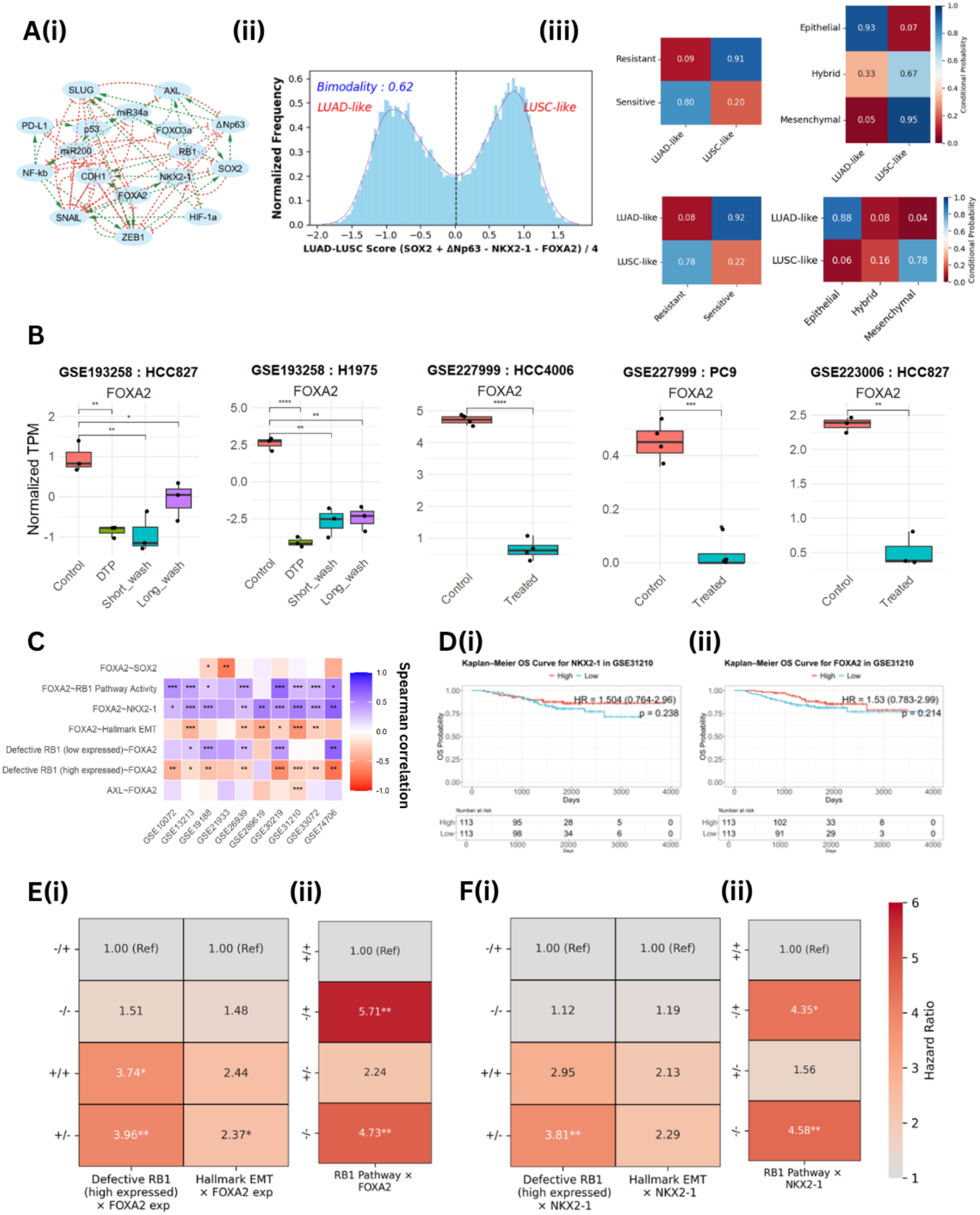
Gene expression and survival analysis for lineage plasticity markers. (A) Modified network to include stemness nodes (SOX2, NKX2-1, FOXA2, ΔNp63) and FOXO3a and HIF-1a; Green colored edges represent activation, and red colored edges represent inhibition. Frequency density histogram of LUAD-to-LUSC score (= (SOX2 + ΔNp63 – NKX2-1 – FOXA2)/4) fitted to Kernel Density Estimation; Conditional probabilities depicting LUAD-Epithelial-Sensitive and LUSC-Mesenchymal-Resistant phenotype likely to occur together. (B) Normalized log2 TPM scores of LUAD-maintaining marker, FOXA2, decrease upon treatment consistently across OSM treatment experiments. (C) Correlation values between normalized FOXA2 expressions with enrichment scores of Hallmark EMT, AXL and RB1 associated signatures show consistent downregulation of FOXA2 with increasing EMT or resistance scores across patient datasets. (D)OS KM-Plots for gene expression levels of (i) FOXA2 and (ii) NKX2-2 in GSE31210, (E) (i, ii) Heatmaps showing Hazard Ratio (HR) values for different combinations FOXA2 gene expression with RB1-pathway activity, Defective RB1 (high expressed) and Hallmark EMT respectively, (F)(i, ii) Same as E (i,ii) but for different combinations: NKX2-1 gene expression with RB1-pathway activity, Defective RB1 (high expressed) and Hallmark EMT respectively (* indicates p-value < 0.05, ** indicates p-value <0.01, *** indicates p-value <0.001; OS: Overall Survival, DFS: Disease Free Survival and PFS: Progression Free Survival, DFS: Disease Free Survival)

We next checked if canonical drivers of LUAD such as FOXA2 are downregulated upon osimertinib treatment. We observed a decrease in FOXA2 expression in DTPs generated from 21-day osimertinib treatment in H1975 and HCC827 cells (GSE193258) (**Fig 7B**). Similar to previous observations about EMT in this dataset, FOXA2 levels were not restored upon short-term or long-term washout of the drug. These trends about FOXA2 downregulation were also witnessed in other independent in vitro datasets: GSE227999 (94) (PC9, HCC4006 cells) and GSE223006 (95) (HCC827 cells). The analysis of multiple patient cohorts exhibited consistent trends for FOXA2 and NKX2-1 expression levels being positively correlated with one another, and positively correlated with RB1 pathway activity scores, but negatively correlated with the Hallmark EMT and with defective RB1 (high expressed) signatures (**Fig 7C, S16D**). Conversely, expression levels for LUSC drivers – SOX2 and ΔNp63 – both correlates positively with defective RB1 (high expressed) in these cohorts (**Fig S15D**). Together, these results indicate that a partial loss of pulmonary lineage identity is a hallmark of osimertinib resistance in EGFR-mutant LUAD.

Next, we interrogated how the expression levels of NKX2-1 or FOXA2 associate with patient survival metrics. In GSE31210, lower levels of either of them associates with worse OS or DFS but not in a statistically significant manner (**Fig 7D, S16E**). In TCGA-LUAD, lower levels of FOXA2 associate with worse OS (HR = 1.26, p = 0.09), DFS (HR = 1.56, p = 0.03), DSS (HR = 1.37, p = 0.07) and PFS (HR = 1.49, p < 0.001) (**Fig 7D, S17A**). We examined whether a decrease in NKX2-1 or FOXA2 – representing a partial lineage switch – along with EMT or altered RB1 activity – led to worse outcomes, as expected based on ‘teams’ like behaviour. We observed that reduced levels of NKX2-1 or FOXA2, along with decreased RB1 pathway activity or with increased scores of Hallmark EMT or defective RB1 (high expressed), led to worse OS and DFS in GSE31210 with HR values up to 4.73 (95% CI = 1.58 - 14.17; p < 0.01) for OS, and up to 4.77 (95% CI = 2.07 - 10.95, p < 0.001) (**Fig 7E-F, S17, S18**). Consistent trends were observed for OS, DFS, DSS and PFS in TCGA-LUAD cohort (**Fig S17, S18**), underscoring that the coordinated simultaneous activation of many axes of plasticity – EMT, partial linage switch from LUAD to LUSC, decreased RB1 pathway activity – lead to worse clinical outcomes than just one axes being activated.

## Discussion

Our study highlights that resistance to EGFR-TKIs, EMT and lineage switch in NSCLC are not merely co-occurring phenomenon but are dynamic processes that can reinforce one another through interconnected feedback loops in the underlying GRN. Such bidirectional regulation, as observed through a) perturbation analysis demonstrating that EMT induction leads to reduced upregulation of RB1 loss genes in HCC827 cells (**Fig 3**), and *vice versa* (9), and knockdown of E-cadherin in HCC827 cells enabling EGFR-TKI resistance, and resistant sublines generated by stepwise escalation leading to spindle-like morphology and altered levels of E-cadherin and Vimentin(96), substantiate our observations that perturbations made along one axis of cell plasticity can trigger coordinated responses along other axes. Overexpression of TWIST1 – an EMT-TF – has been shown to be necessary and sufficient for resistance to EGFR-TKIs in EGFR-mutant NSCLC *in vitro* and *in vivo* in a subset of cell lines (97).Similar reports indicate the role of SLUG and ZEB1, both of which can repress E-cadherin along with TWIST1 (98,99). Therefore, inhibiting EMT through mechanisms such as differentiation therapy can be a viable therapeutic strategy.

Besides EGFR-TKI resistance, EMT-associated molecules have been identified as common mechanisms of primary and/or acquired resistance to targeted therapy – ALK inhibitors (100,101) and KRAS G12C inhibitors (102,103) in lung cancer, tamoxifen in ER+ breast cancer (104,105) and enzalutamide resistance (106) in prostate cancer. These observations support the emerging notion that higher plasticity unlocked through at least a partial activation of EMT may enable cells under therapeutic assault to explore new ‘attractors’ in their high-dimensional phenotypic space, enabling adaptive cell-state transitions (107). Higher plasticity is a hallmark trait of hybrid epithelial/ mesenchymal (E/M) phenotypes (31,108) that have also been reported to emerge in *in situ* long-term osimertinib resistance assays (109). Thus, an indirect selection for higher phenotypic plasticity and enhanced evolvability can be amplified as cells adapt to various orthogonal therapeutic stressors, eventually leading to increased metastatic competence (110).

PD-L1 is a central immune checkpoint molecule that enables tumour immune evasion and is widely used as a predictive biomarker for immunotherapy response. Previous studies have demonstrated that AXL expression is positively correlated with PD-L1 levels, and that elevated AXL expression in PD-L1–high tumours can influence clinical outcomes following immunotherapy (111,112). PD-L1 expression consistently co-occurs with high AXL expression and mesenchymal markers in patient bulk transcriptomic datasets. In contrast, RB1 loss–associated signatures do not display consistent relationships with PD-L1 across bulk, single-cell, and spatial datasets. At single-cell resolution, RB1 pathway activity shows a negative correlation with PD-L1 signatures consistent with the opposing “teams” predicted by our network model. The discrepancy between bulk and single-cell observations likely arises from the strong influence of immune infiltration and tumour microenvironment on PD-L1 expression in bulk profiles. These confounding effects are minimized at single-cell and spatial resolutions, thereby revealing weak negative association between RB1 pathway activity and PD-L1 signature (**Fig S3, S5A)**.

We focused on LUAD to LUSC transition; however, NSCLC to SCLC transformation has also been observed in EGFR-TKI and ALK-TKI resistance (113–115). It remains unclear yet whether these two trajectories of lineage plasticity are mutually exclusive or not, i.e. whether cells transitioning to SCLC necessarily navigate through LUSC. Similarities and differences in molecular profiling of these two scenarios at a multi-omics level would be critical in mapping the therapeutic vulnerabilities for both these cases. We observed an association between EMT and basal-shift transformation to LUSC, but the involvement of EMT in transformed SCLC is likely to be more diffused, given the heterogeneity in SCLC along the neuroendocrine/non-neuroendocrine (NE/non-NE) spectrum (116,117). Future efforts focusing on decoding necessary and sufficient conditions at cell-intrinsic and micro-environmental levels shaping these phenotypic plasticity landscapes would be crucial.

Our findings have several therapeutic implications. First, they suggest that osimertinib resistance is maintained by a coordinated plasticity network rather than by a single linear bypass pathway, arguing against reliance on one-dimensional biomarker strategies alone. The recurrent coupling of EMT, AXL, PD-L1, and reduced RB1 activity supports rational co-targeting approaches aimed at destabilizing the mesenchymal/resistant state. In this context, combinations of EGFR inhibition with agents that suppress AXL signalling, EMT-associated programs, or inflammatory/NF-κB signalling may be more effective than sequential monotherapies in delaying or preventing adaptive escape. Second, the observation that resistant states can persist after drug withdrawal is consistent with cellular memory and suggests that early intervention against plasticity programs may be more effective than attempting to reverse fully established resistant states. Third, the association of combined plasticity programs with worse survival indicates that composite signatures integrating EMT, AXL, PD-L1, and RB1 pathway activity may provide greater prognostic and therapeutic value than any single marker alone. More broadly, our results support a treatment paradigm in which the goal is not only to inhibit the dominant oncogenic driver, but also to collapse the network architecture that enables adaptive cell-state transitions under therapeutic stress. Such topology-informed strategies may help identify patients at high risk for adaptive resistance and guide combination therapies designed to restrict tumour evolvability before overt progression emerges.

Our study has certain limitations. First, we have focused mostly on transcriptional layer of regulatory control, without performing an integrative analysis of crosstalk at metabolic (118) and/or epigenetic levels (119). Second, we do not explicitly consider changes in the tumour microenvironment during osimertinib resistance, instead focus on cancer cell-autonomous reprogramming (120). Third, our model focuses on mapping the steady-state behaviour of underlying GRNs. Despite these limitations, our findings provide molecular underpinnings for how EMT, lineage plasticity and AXL-RB1 pathway driven osimertinib resistance can reinforce one another, unlocking coordinated plasticity along multiple functional axes due to the presence of ‘teams’ of nodes in the underlying GRNs. It offers a framework to understand why recent efforts such as bispecific antibody that targets EGFR and AXL can delay tumour relapse and views the emergence of resistance as a dynamic evolution process instead of a static molecular tag (121). It also underscores why single-point disruptions are likely to be ineffective, and future therapeutic designs needs a conceptual shift to attempt network topology based intelligent perturbation.

## Acknowledgements

MKJ was supported by Param Hansa Philanthropies. AT was supported by the NIH Intramural Research Program (NCI ZIA BC 011793). The contributions of the NIH author(s) were made as part of their official duties as NIH federal employees, are in compliance with agency policy requirements, and are considered Works of the United States Government. However, the findings and conclusions presented in this article are those of the author(s) and do not necessarily reflect the views of the NIH or the U.S. Department of Health and Human Services.

## Author contributions

MKJ conceived and supervised research and obtained funding. SV and RKM performed research and wrote the first draft of the manuscript. All authors contributed to data analysis and editing of the manuscript.

## Data availability

All codes for the manuscript are available at: https://github.com/Shreya0217/Osm_res_EGFR_LUAD.git

## Methods

### Network simulations

#### RACIPE

RACIPE framework was applied for continuous dynamic simulations of Gene Regulatory Networks (GRNs) (41). RACIPE samples parameters and random initial conditions over a specified range resulting in possible steady-state behaviors of the network.

A topology file for the network containing information about activation and inhibition links between nodes was created and number of parameter sets was set to 10,000. RACIPE samples multiple initial parameters from the designated range for each parameter and initial conditions from a log-uniform distribution spanning the minimum to maximum expression levels for that node. RACIPE solves shifted Hill-function based ordinary differential equations (Equation 1) numerically to give possible steady states as outputs.

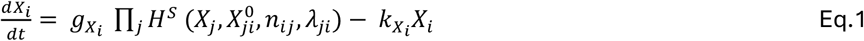

Here, *X*_*i*_ denotes the _*i*_ concentration of the gene product corresponding to _*i*_ node *X*_*i*_ in the gene regulatory network _*ij*_ (GRN). The term 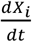 represents the rate of change of gene expression over time. The parameter 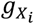 indicates the basal production rate, while 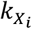 denotes the basal degradation rate of the gene product *X*_*i*_. The term *H*^*s*^ refers to the shifted Hill function, where *n*_*ij*_ represents the Hill coefficient, 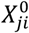 denotes the threshold value, and *X*_*j*_ represents the concentration of the gene product corresponding to node *X*_*i*_. Here, *X*_*j*_ is a node that either activates or inhibits *X*_*i*_.

RACIPE produced *log*_2_-normalized steady-state gene expression levels as output, which was subsequently z-normalized for further downstream analysis.

### Epithelial-Mesenchymal (EM) and resistance scores

EM score and resistance score for each steady state was calculated as difference of expression values of certain nodes as defined in Equation 2 and 3.

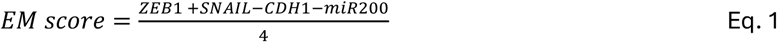

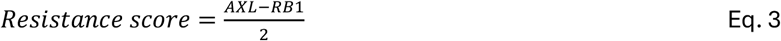

Histogram of these score vectors was plotted and fitted with Gaussian Mixture model (GMM) to cluster the states to specify phenotypes and resistance status. Principal Component Analysis (PCA) was performed on the z-normalized steady-state gene expression values, and the scatter points were color coded with phenotypes derived from GMM clusters of scores. Conditional probabilities were calculated of EM phenotype given a resistance status and vice versa based on frequency of steady states. The bimodality in resistance scores is quantified by calculating Sarle’s bimodality coefficient (Equation 4), where *γ* is skewness, *κ* is kurtosis and n is the number of observations in a distribution (122).

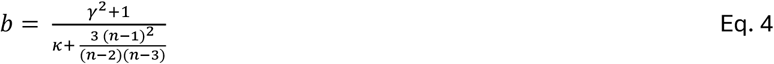

### Team strength analysis

#### Adjacency and Influence Matrix

The team strength calculations were based on previous study from our group (31). Adjacency matrix for the input network topology was defined as given in Equation 5.

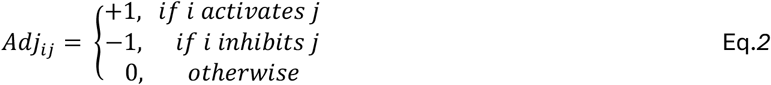

The adjacency matrix values are set as +1 or -1 only if there is direct inhibition or activation respectively and 0 when there is not direct edge between nodes. To capture the interactions between nodes over longer path lengths, Influence matrix, *Inf l* has been defined in Equation 6.

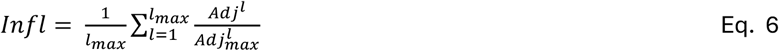

Here, *Adj*^*l*^ represents the adjacency matrix raised to the power *l*, while 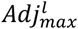 denotes the adjacency matrix in which all inhibitory edges are replaced with activating edges. The term 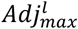 represents element-wise division between the corresponding entries of the two matrices. The summation over *l* is normalized by *l*_*max*_ to constrain the values of the influence matrix within the range [*−*1, 1], thereby preventing numerical blow-up.

#### Identification of “teams”

A set of nodes *N* is defined to be in team *T* if *Infl*_*ij*_*>* 0 for all *i,j* ∈ *T* . The two teams in this study are identified using hierarchical clustering on the matrix *M*, where *M* is constructed by horizontally concatenating the influence matrix with its transpose. This matrix captures both incoming and outgoing interactions for each node.

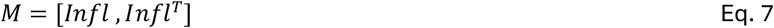

The team strength of the network is defined as:

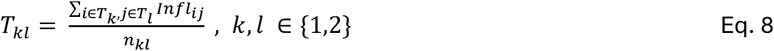

Here, *n*_*kl*_ denotes the number of interactions between teams *T*_*k*_ and *T*_*l*_. This formulation yields four quantities: *T*_11_, *T*_12_, *T*_21_ and *T*_22_ representing intra-team and inter-team interactions.

The overall team strength of the network is defined as the average of the absolute values of these terms:

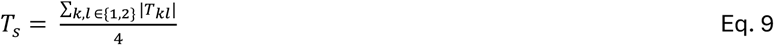

### Random Network Analysis

The random network analysis was conducted to understand uniqueness of biological wild-type (WT) network as compared to non-biological networks. These non-biological networks are generated by random shuffling of edges in the WT network, followed by RACIPE simulations and team strength calculations for each network.

### Gene expression data analysis

Publicly available bulk RNA-seq datasets for EGFR-mutant NSCLC cell lines treated with OSM were obtained from GEO. Lung adenocarcinoma (LUAD) and lung squamous cell carcinoma (LUSC) patient datasets were obtained from TCGA, with additional patient cohorts retrieved from GEO. RNA-seq expression data were analyzed using TPM-normalized expression values. Pre-processed RMA-normalized Affymetrix microarray datasets were taken by using getGEO from GEOquery R package (http://bioconductor.org/packages/GEOquery/). Expression datasets for lung cancer cell lines were obtained from the Cancer Cell Line Encyclopedia (CCLE) (45). Single-sample gene set enrichment analysis (ssGSEA) for the different gene signatures was performed using the GSVA R package (46,124).

Since the genes in RB1 Pathway Activity represent downregulated genes upon RB1 activation, the enrichment scores were inverted by taking negative of them and scaling them across samples. KS score was calculated using 2KS test statistics described in (48).

### AXL signature

To derive an AXL gene signature, we found top correlated genes with AXL across TCGA-LUAD and TCGA-LUSC dataset. For each dataset, genes showing a strong and significant positive association with AXL expression (Spearman correlation > 0.5, p-value < 0.01) were selected.

### Single cell RNA sequence analysis

Publicly available single-cell RNA-seq datasets were obtained from GEO; detailed descriptions of all datasets are provided in Supplementary Table X. Raw count matrices were converted to TPM-normalized expression values prior to downstream analysis. The TPM-normalized feature barcode matrices were imputed using Markov Affinity-based Graph Imputation of Cells (MAGIC) to mitigate dropout effects using Rmagic R package (125). Custom filtering was performed for each dataset and manual annotation of cells based on epithelial, stromal and immune markers are performed. Gene set activity was quantified using an area under the recovery curve (AUC)–based approach, which measures the enrichment of gene set members within the ranked gene expression profile of each cell. AUC scores for each signature were computed using the AUCell R package (73). For OSM-treated samples, box plots were constructed using the mean AUC score across all cells within each sample. All single-cell datasets were handled in R using the Seurat package (126). Again, for RB1 Pathway Activity, the final scores were calculated by subtracting AUC scores from maximum of AUC across all cells.

### Spatial transcriptomic data analysis

One patient sample each from two publicly available NSCLC patient spatial transcriptomics datasets were used to perform spatial transcriptomics analysis. 10X Visium datasets were obtained from GEO for GSE189487 and EMBL-EBI for E-MTAB-13530 and activity scores for different gene sets were calculated using AUCell. All analyses were using R package Seurat.

### Survival analysis

For signature scoring, ssGSEA was performed using GSVA package (124). For each signature, samples were classified into “+” (high) and “-” low groups based on the cohort-specific median of the ssGSEA score. For Cox-hazard ratio analysis, we performed three levels of grouping models. Groups were defined as follows: one-axis (single signature) plotted as Kaplan Meier plots, two-axis (pairwise combinations) plotted as heatmaps, and three-axis (triple combinations) plotted as heatmaps. The KM survival curves were generated using the ggsurvplot function in survminer package in R (126). Cox proportional-hazard models were fitted using the survival package with reference groups defined a priori. Heatmaps of hazard ratios and 95% confidence intervals were generated using the survminer package.

## List of Supplementary Tables

| S.No | Table description |
| --- | --- |
| 1 | Table 1: References for network edges for Fig 1A and Fig S2 |
| 2 | Table 2: References for network edges for Fig 7A |
| 3 | Table 3: Description of publicly available perturbation in cell line datasets from Gene Expression Omnibus (GEO) |
| 4 | Table 4: Description of publicly available patient datasets from Gene Expression Omnibus (GEO) |

## Supplementary figures

**Figure S 1.**
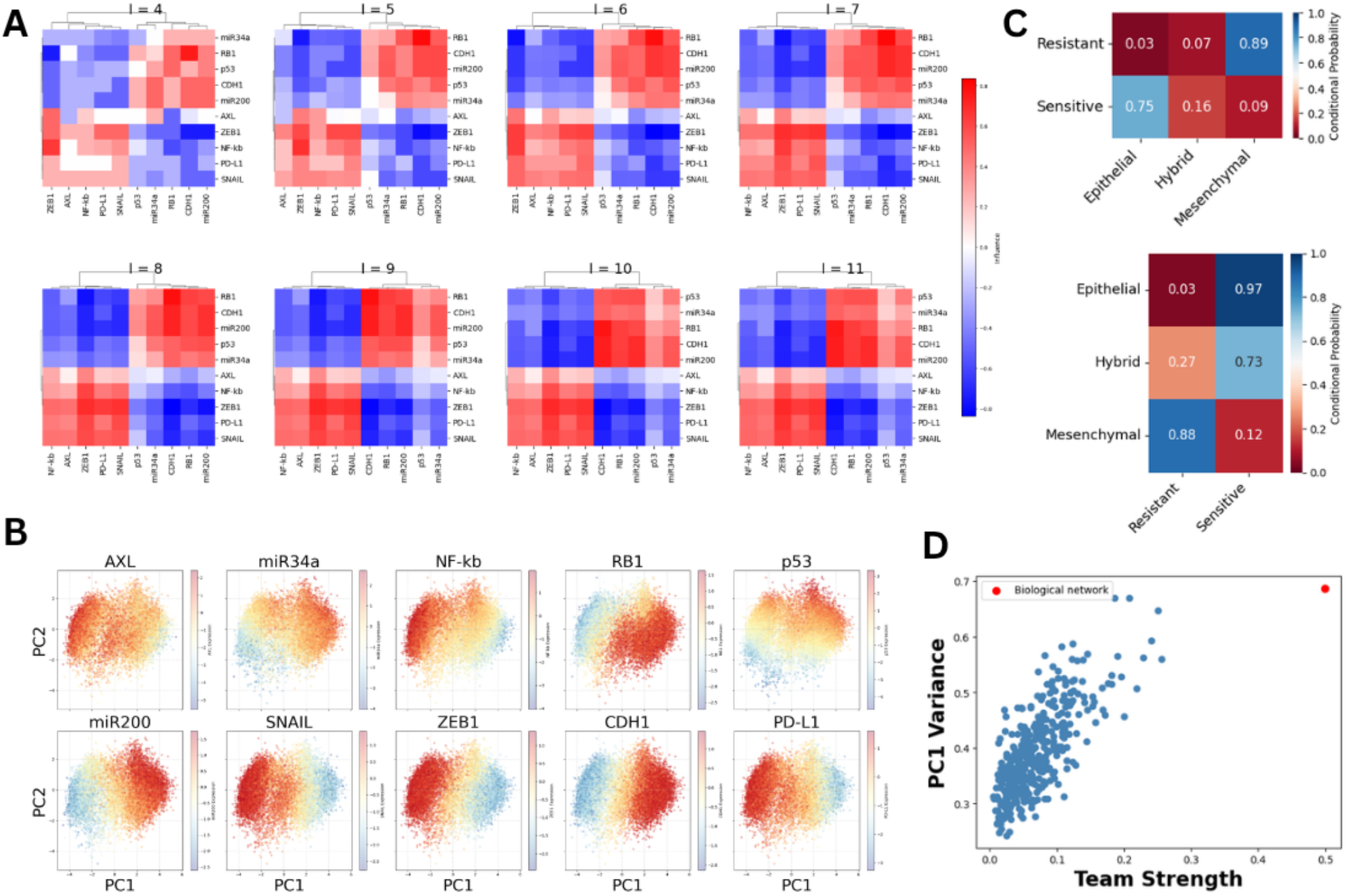
Dynamical simulations of EMT-RB1-AXL gene regulatory network (GRN). (A) Influence matrix evolution with increasing maximum path length. (B) PCA dimensionality reduction plots color coded by network nodes steady state values. (C) Conditional probabilities of resistance status (rows) given EM phenotype (columns) and vice versa. (D) Scatter plot of team strength and PC1 variance showing biological network with highest team strength and PC1 variance.

**Figure S2.**
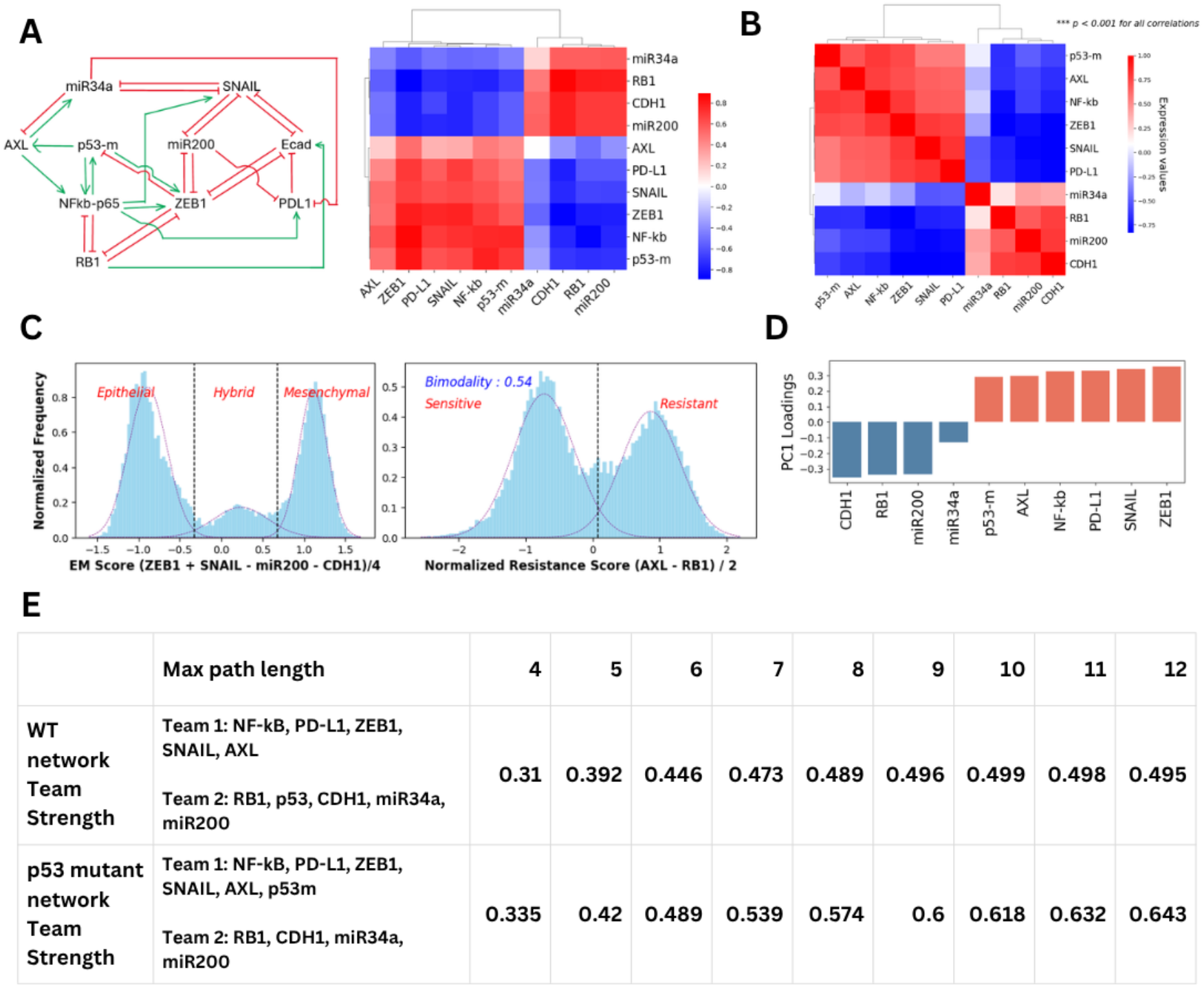
Dynamical simulations of the p53 mutant version of the gene regulatory network. (A) Modified network architecture representing p53 gain-of-function (GOF) mutation. Influence matrix of the network computed with a maximum path length of 10. (B) Density histograms of EMT score (= (mir200 + Ecad − ZEB1 − SNAIL)/4) and resistance score (= (AXL − RB1)/2), fitted to Gaussian mixture models with three and two components, respectively. (C) Principal component (PC1) loadings for each node showing p53-mutant node flipping teams. (E) Team strength values for different path lengths for wild type p53 and mutant p53 networks.

**Figure S3.**
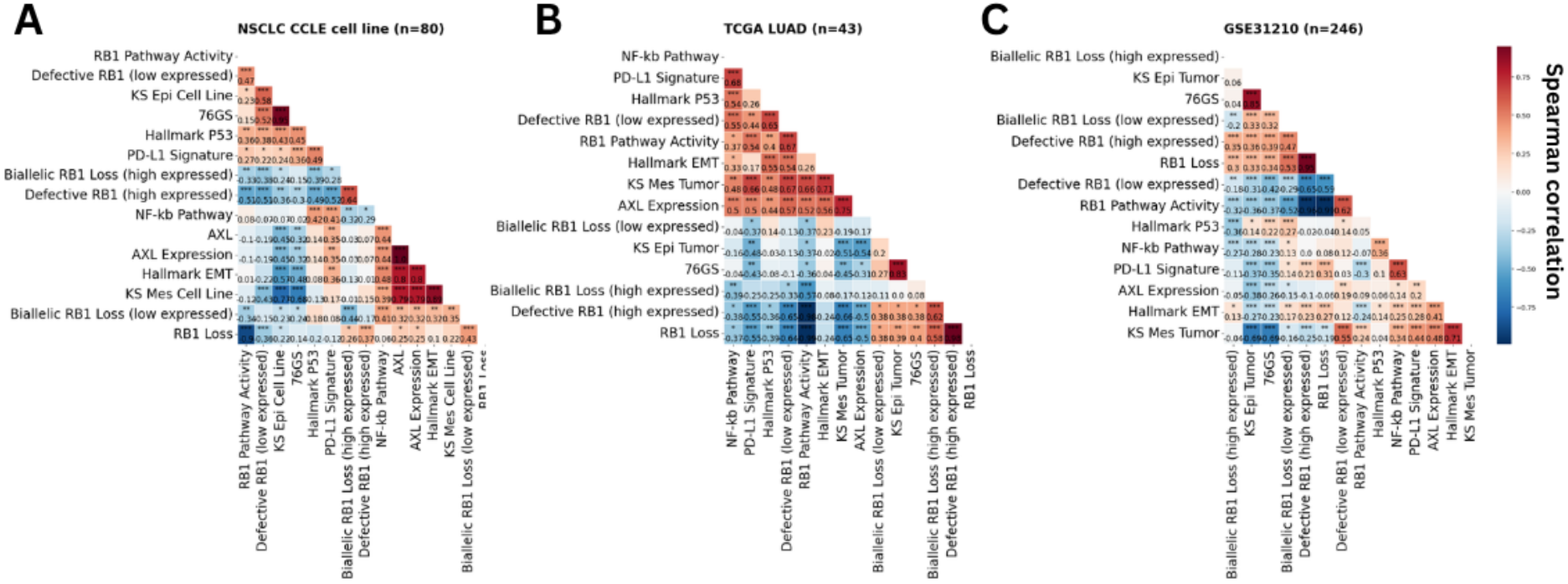
**Correlation plots of ssGSEA scores of different signatures in** (A) 80 NSCLC cell lines from CCLE database. (B) 44 EGFR mutant NSCLC samples from TCGA database. (C) GSE33072 dataset containing residual tumor samples from first generation EGFR TKI drug treatment from BATTLE trial.

**Figure S4.**
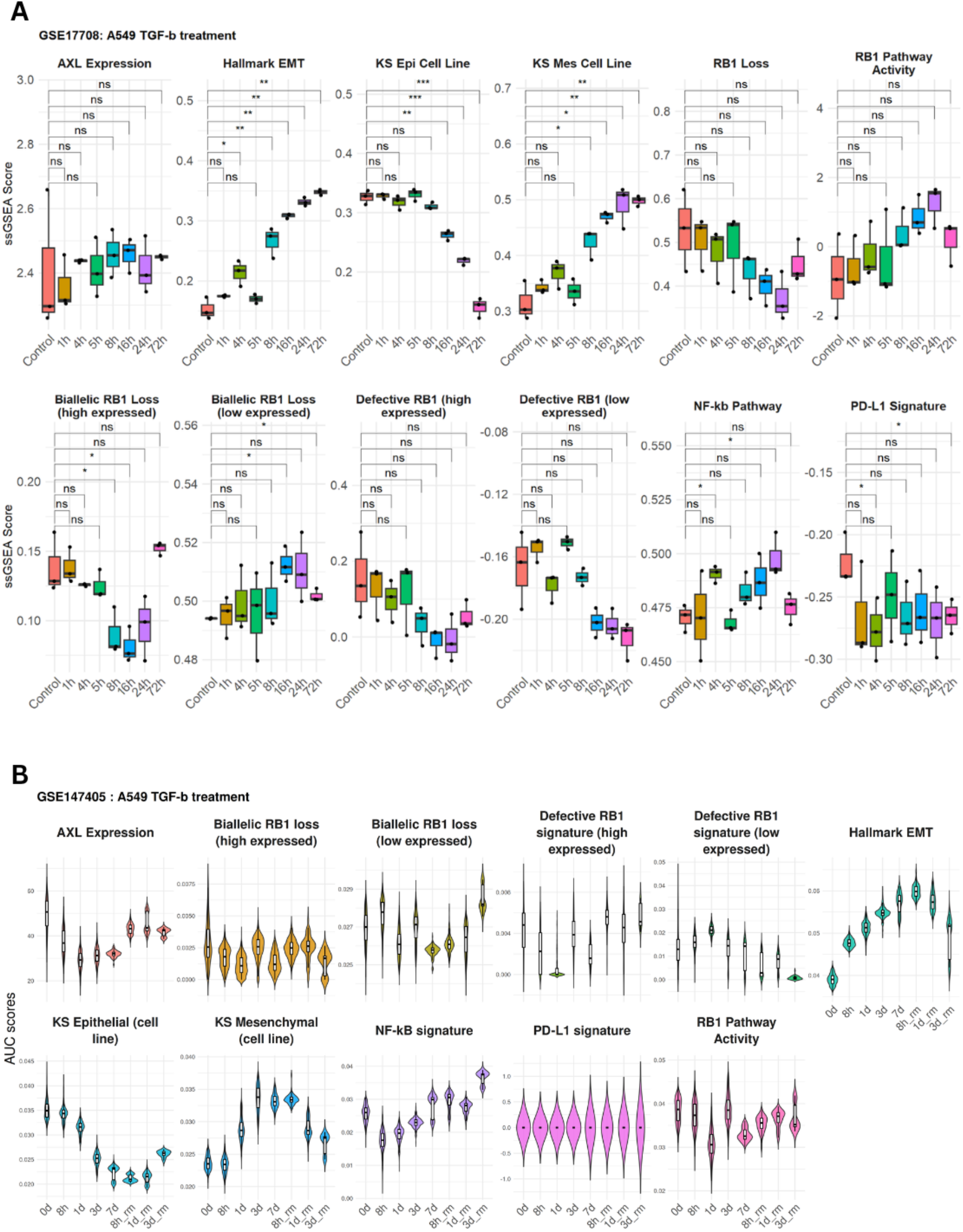
Enrichment scores in time course EMT induction in KRAS-mutant cell lines A549 depicting non-significant changes with respect to control for resistance determining signatures.

**Figure S5.**
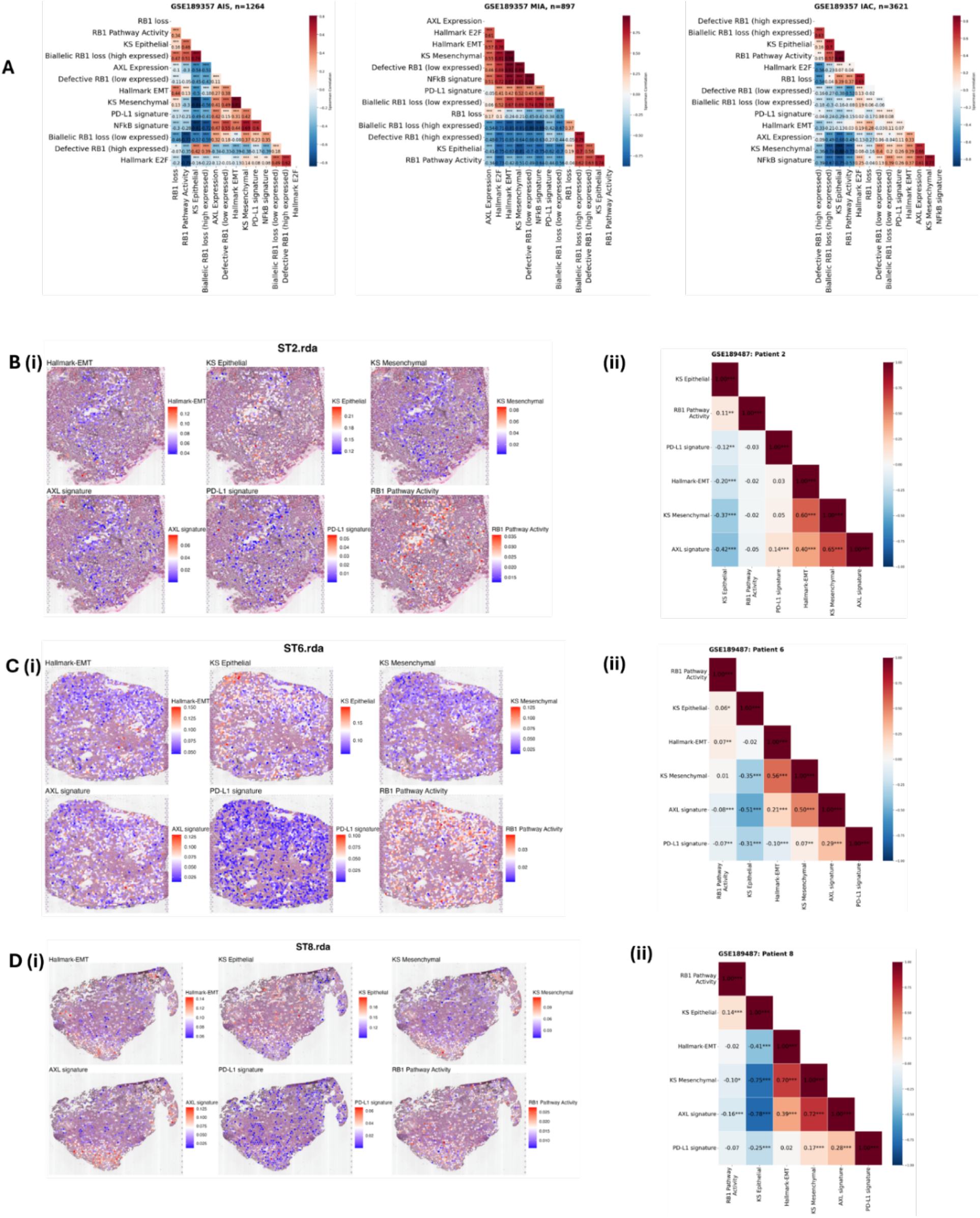
Single cell and spatial transcriptomic analysis for patients and cell lines. (A) Correlation heatmap for different stages of LUAD from AIS to IAC depicting correlation of AUC scores for all gene signatures (GSE189357). (B)(C)(D) Spatial distribution plots for activity of (i) Hallmark-EMT, (ii) KS-Epithelial (iii) KS-Mesenchymal (iv) AXL-signature (v) RB1-pathway activity signatures and pairwise correlation heatmap for activity levels in GSE189487.

**Figure S6.**
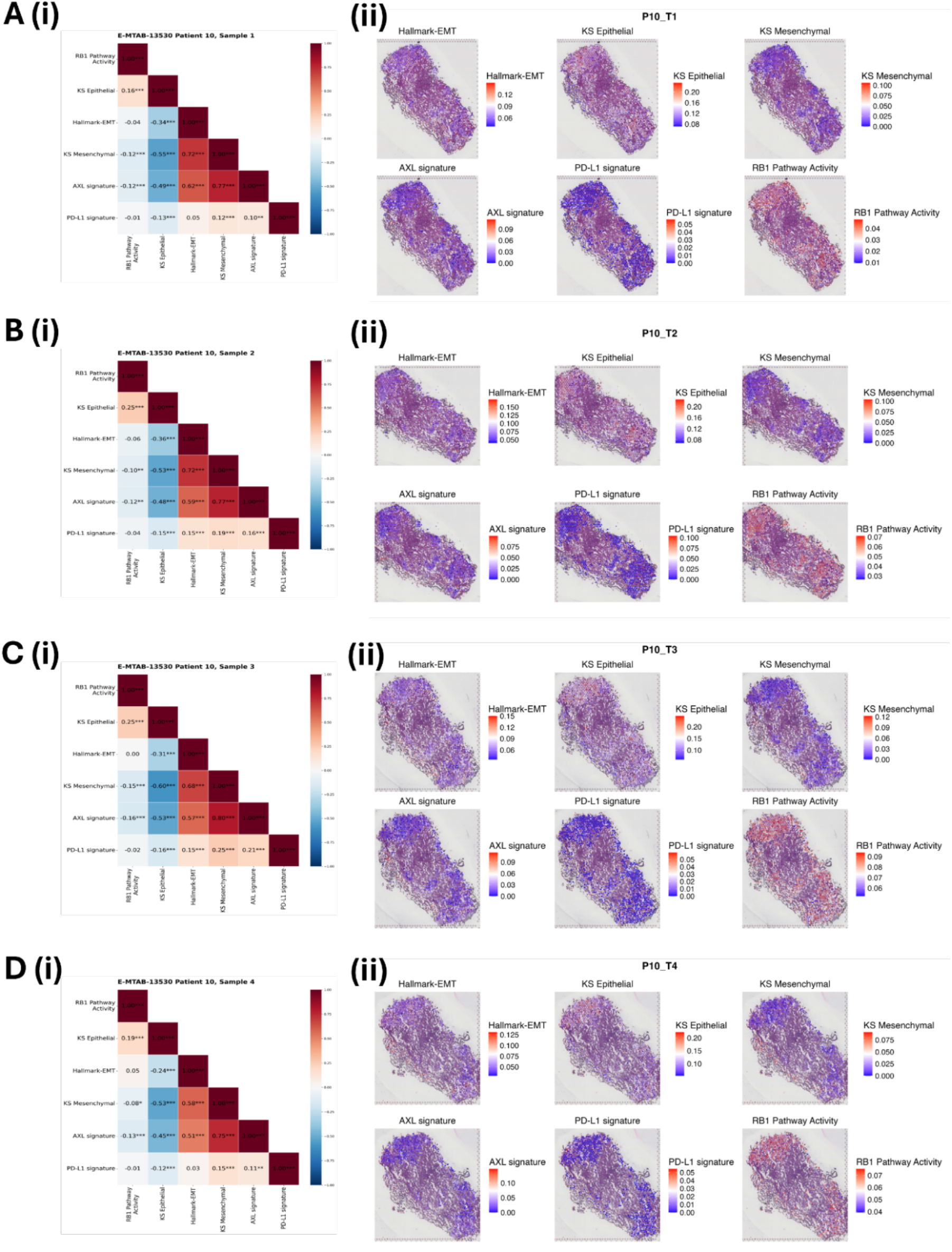
Spatial transcriptomic analyses of four different tumor samples from one patient from E-MTAB-13530.

**Figure S7.**
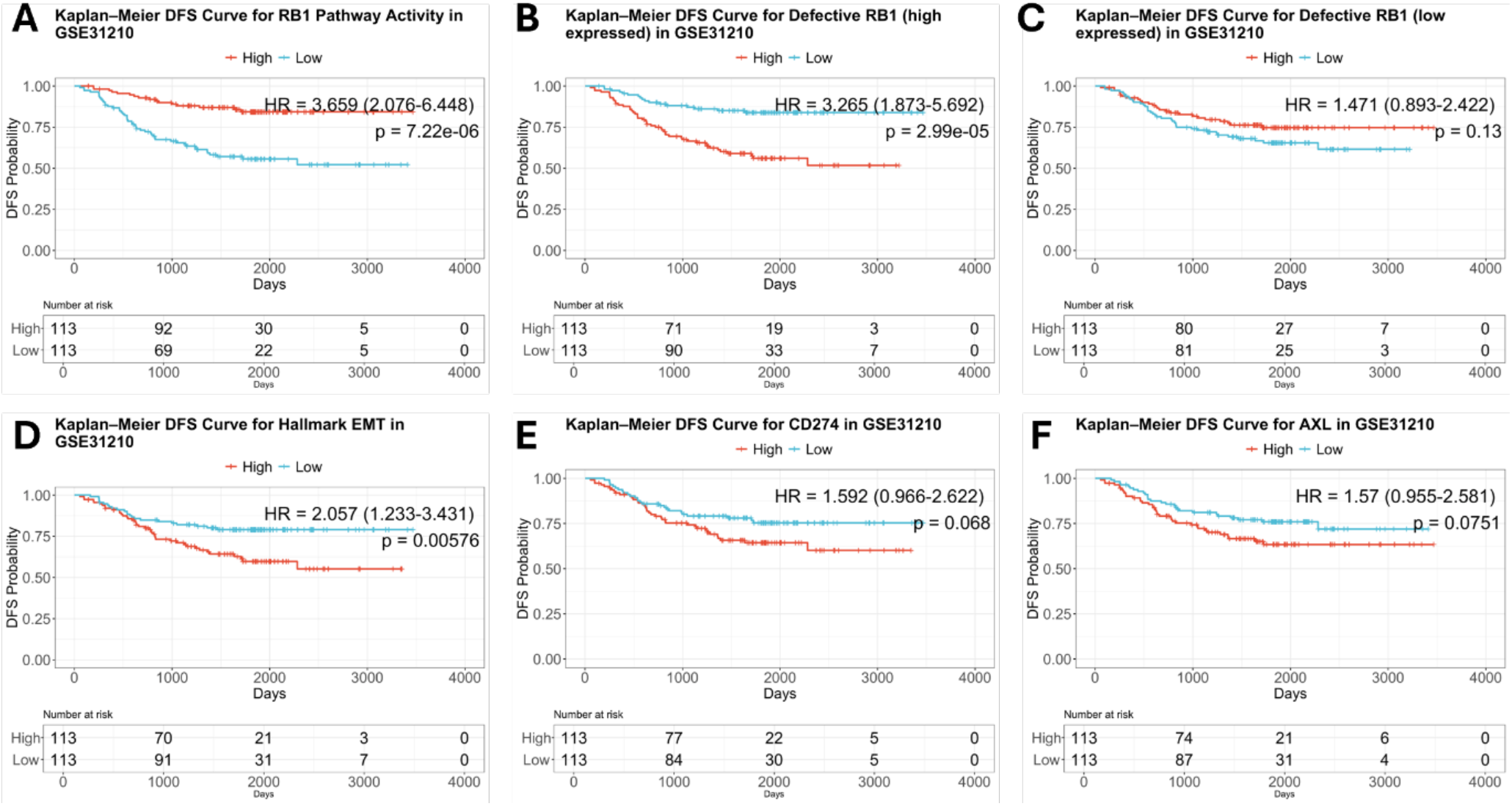
Disease-Free Survival Kaplan-Meier (KM) plots for enrichment scores of different gene sets and expression levels of genes in GSE31210 patient cohort separated by median. (A) Disease Free Survival KM plots for activity levels of Hallmark-EMT gene set (B, C) Disease Free Survival KM plots for expression levels of CD274 (PD-L1 gene) and AXL genes (D, E) Disease Free Survival KM plots for activity for levels of Defective-RB1 (high expressed) and Defective-RB1 (low expressed) gene sets. (DFS: Disease-Free Survival)

**Figure S8.**
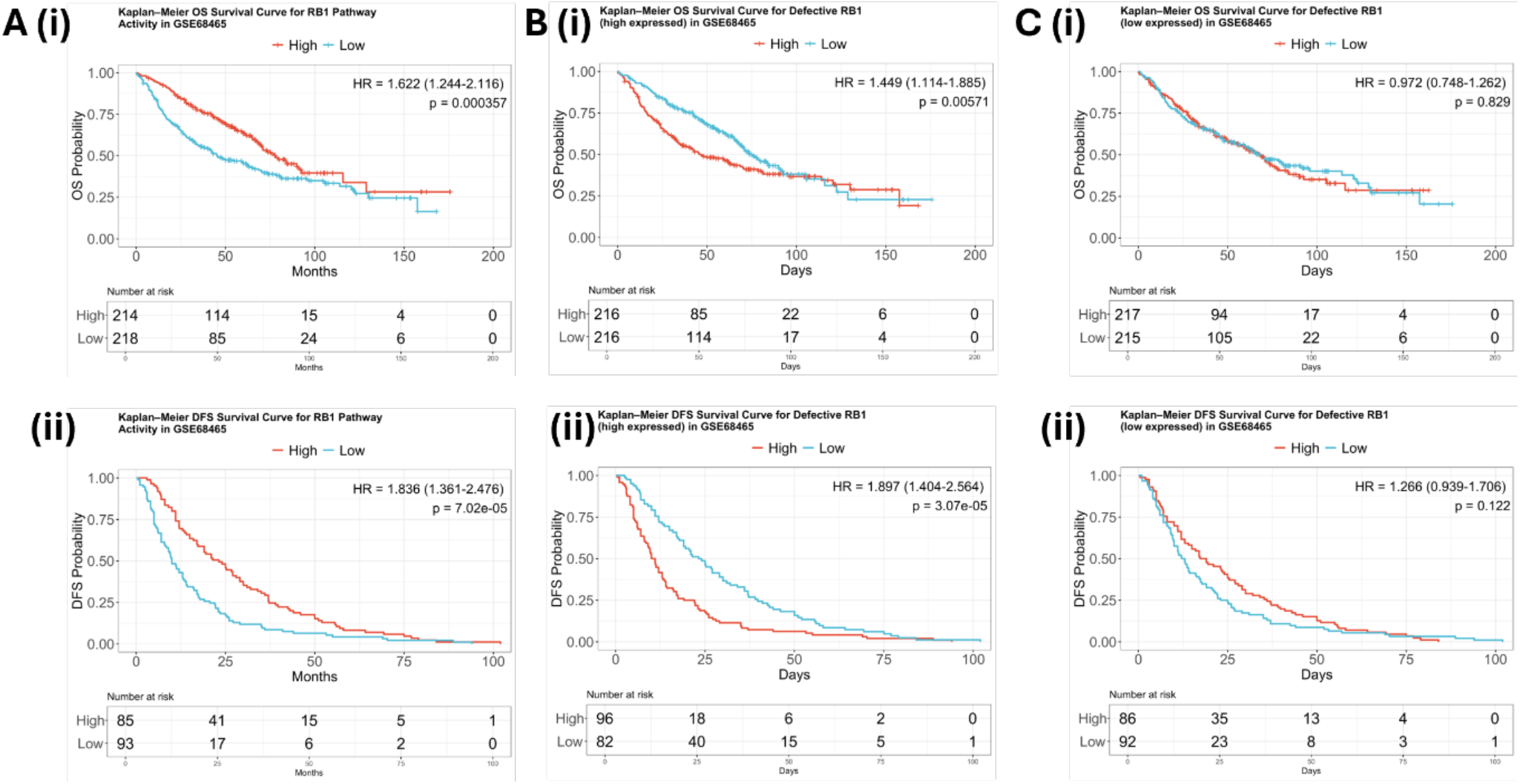
Overall Survival and Disease-Free Survival Kaplan-Meier (KM) plots for enrichment scores of different gene-sets and expression levels of genes in GSE68465 patient cohort separated by median. (A) (i)Overall Survival and (ii) Disease-Free survival KM plots for Defective RB1 loss (high expressed) (B) (i) Overall Survival and (ii) Disease Free Survival KM plots for Defective RB1(low expressed). (OS: Overall Survival; DFS: Disease Free Survival)

**Figure S9.**
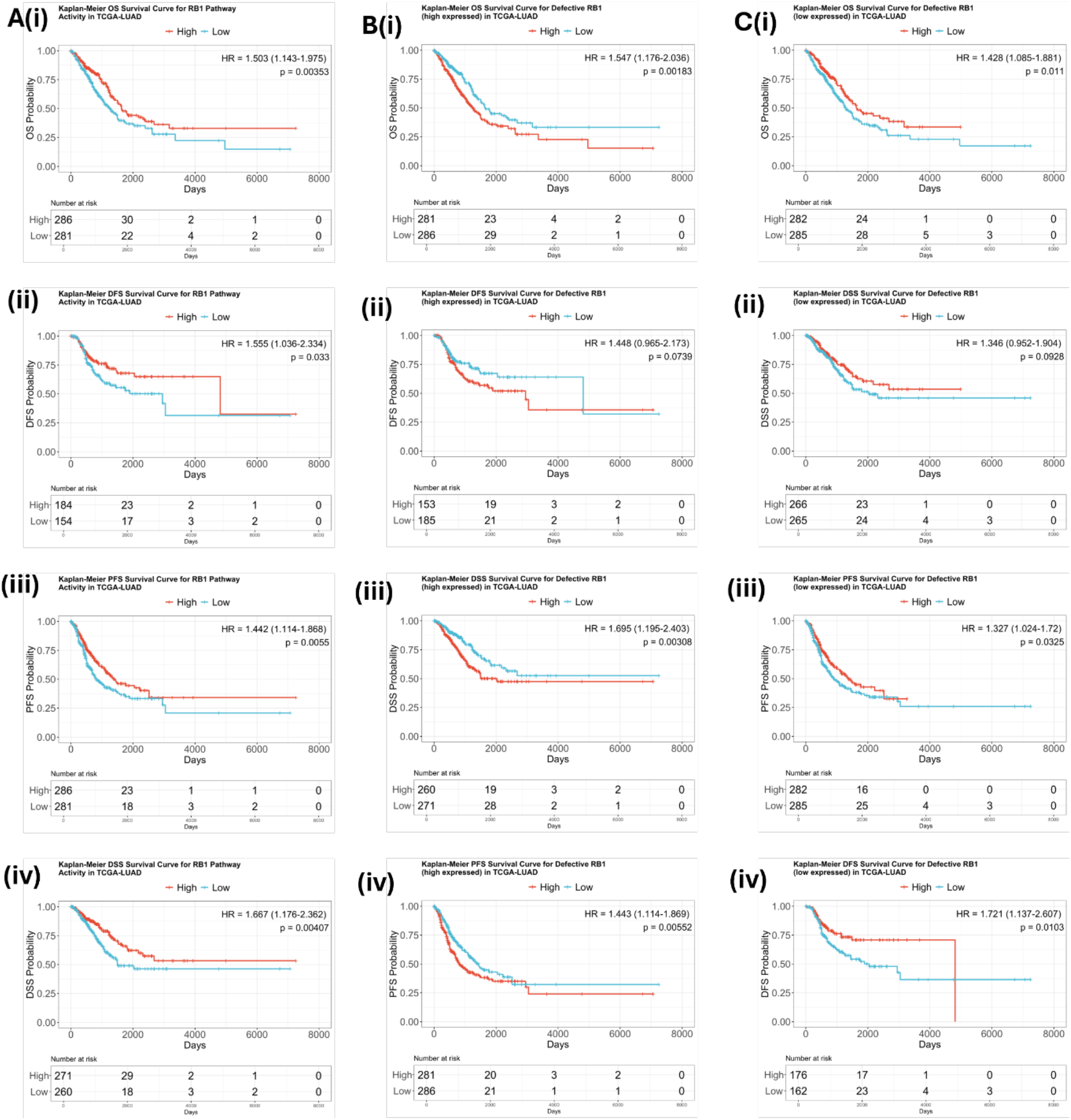
Overall survival (OS), Disease-Free Survival (DFS), Progression Free Survival (PFS), Disease Specific Survival (DSS) Kaplan-Meier (KM) plots for enrichment of different gene sets and genes in TCGA-LUAD patient cohort separated by median. (A) Kaplan-Meier (KM) plots for (i) OS, (ii) DFS, (iii) PFS and (iv) DSS for Defective RB1 loss (high expressed) gene set; (B) Kaplan-Meier (KM) plots for (i) OS, (ii) DFS, (iii) PFS and (iv) DSS for Defective RB1 loss (low expressed) gene set. (OS: Overall Survival; DFS: Disease-Free Survival; PFS: Progression Free Survival; DSS: Disease Specific Survival)

**Figure S10.**
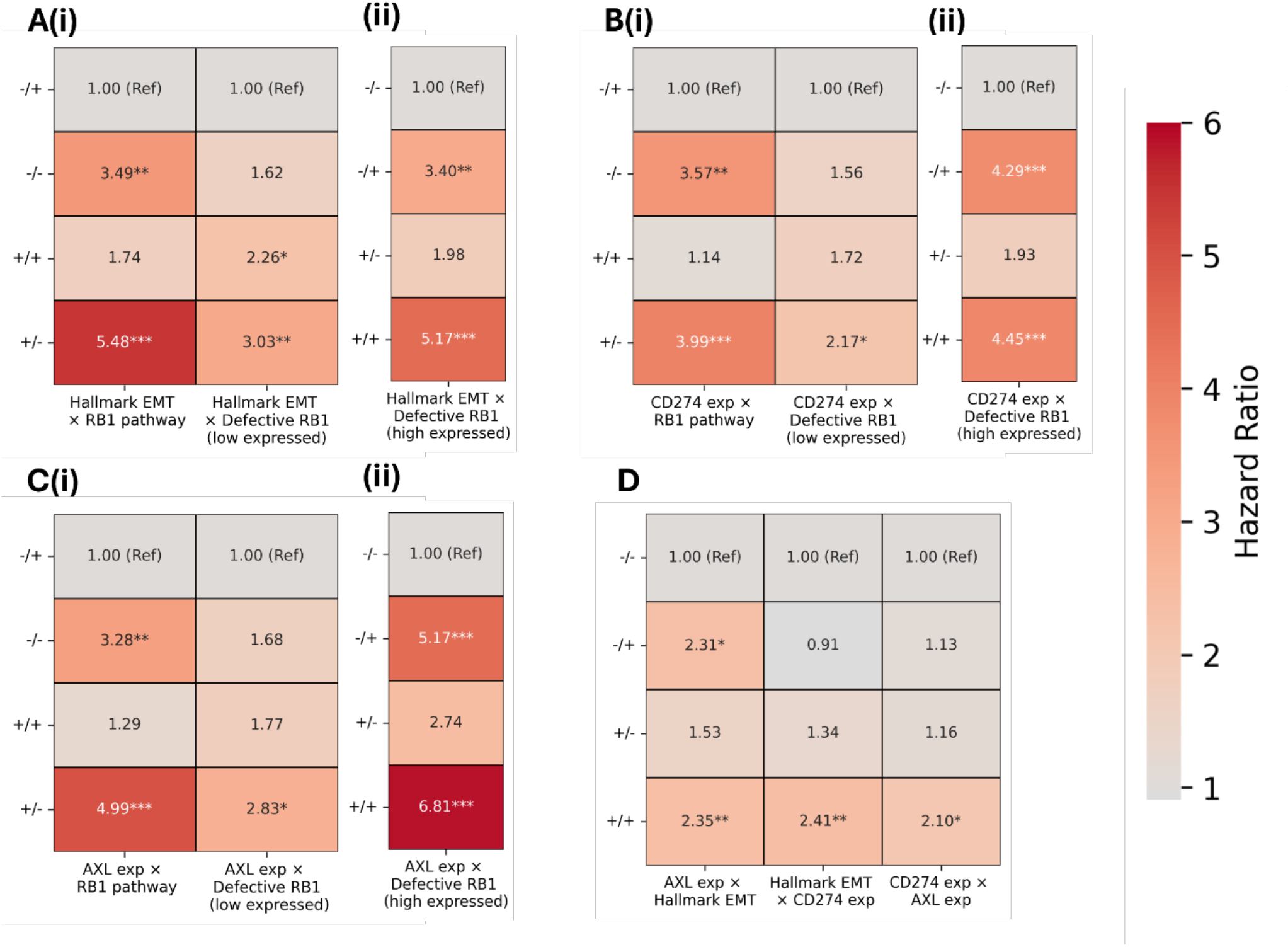
Heatmaps of Disease Free Survival (DFS) Hazard Ratio (HR) values for double combinations of enrichment scores of different gene sets and expression levels of genes in GSE31210. (A) DFS HR values for different combinations of Hallmark-EMT vs (i) RB1 Pathway, Defective-RB1-loss (low expressed) and (ii) Defective-RB1-loss (high expressed) (B) Same as (A) for different combinations of CD274 (PD-L1 gene) vs (i) RB1 Pathway, Defective-RB1-loss (low expressed) and (ii) Defective-RB1-loss (high expressed) (C) Same as (A) for different combinations of AXL gene vs (i) RB1 Pathway, Defective-RB1-loss (low expressed) and (ii) Defective-RB1 loss (high expressed) (D) Same as (A) for different combinations of AXL gene vs Hallmark-EMT, Hallmark-EMT vs CD274 (PD-L1 gene), CD274 (PD-L1 gene) vs AXL gene (* p < 0.05, ** p < 0.01, *** p < 0.001; OS: overall survival; DFS: disease-free survival; PFS: progression-free survival) )

**Figure S11.**
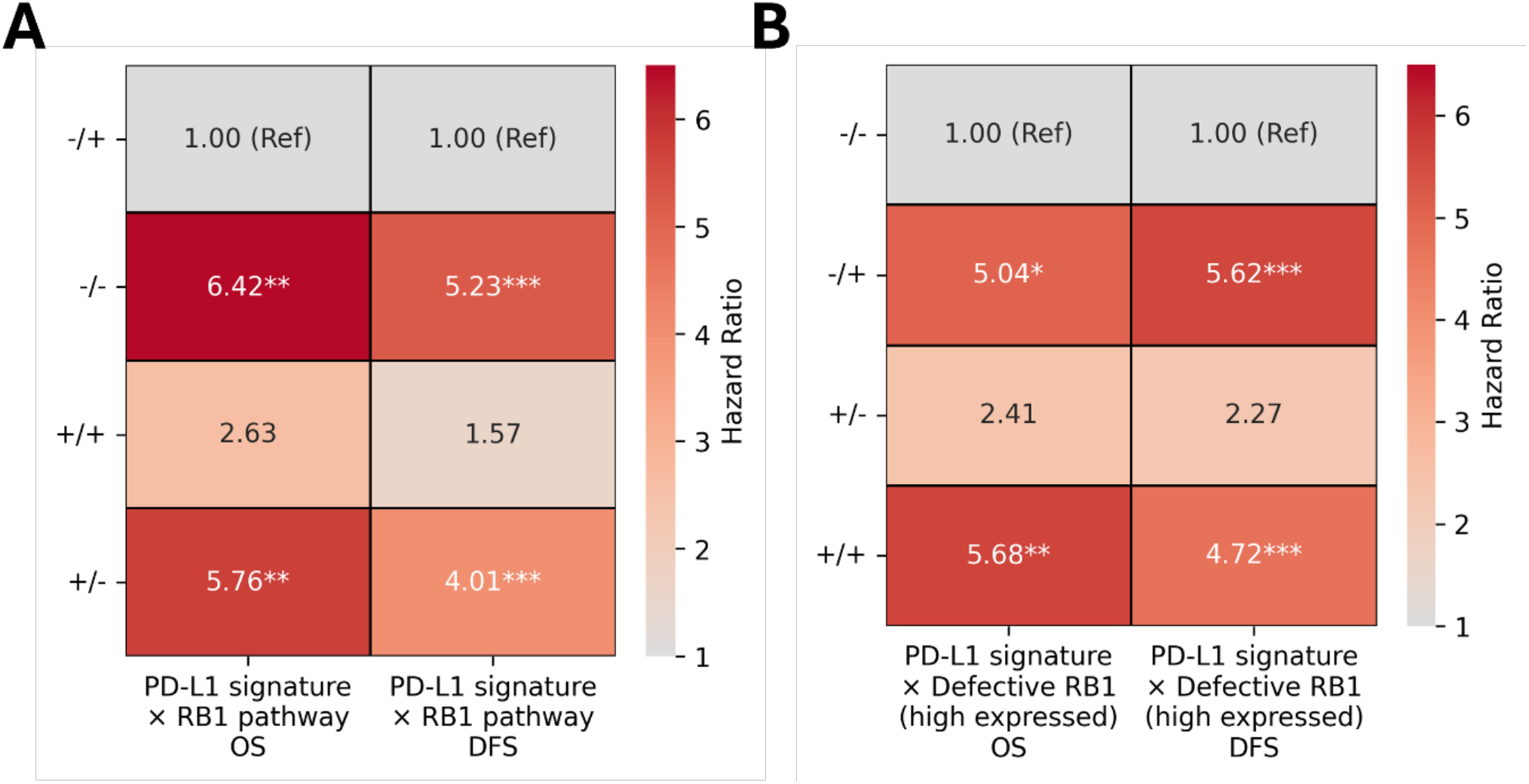
Overall Survival (OS) and Disease-Free Survival (DFS) heatmaps depicting Hazard Ratio (HR) values for double combinations of enrichment scores of different gene sets with PD-L1 signature in GSE31210. (A) OS and DFS HR values for combinations of PD-L1 signature vs RB1-pathway gene sets; (B) Same as (A) for combinations of PD-L1 signature vs Defective-RB1 (high expressed) gene sets. (* indicates p-value < 0.05, ** indicates p-value <0.01, *** indicates p-value <0.001; OS: Overall Survival, DFS: Disease Free Survival)

**Figure S12.**
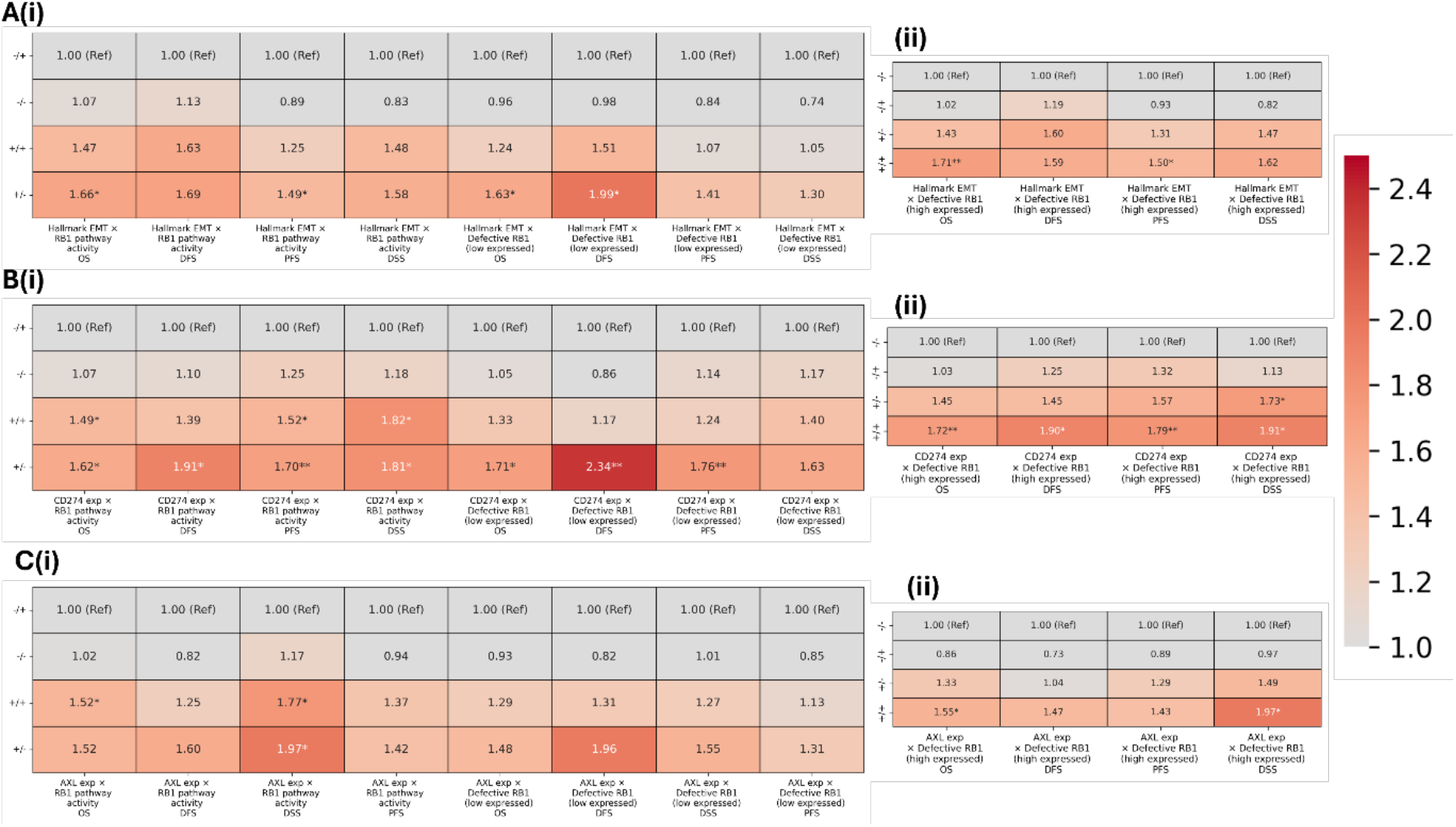
Heatmaps depicting Hazard Ratio (HR) values for PFS, OS, DSS, DFS for double combinations of enrichment scores of different gene-sets and expression levels of genes in TCGA-LUAD dataset with patients separated by median. (A) OS, DFS, PFS and DSS HR values for different combinations of Hallmark-EMT vs (i) RB1 Pathway Activity, Defective-RB1-loss (low expressed) and (ii) Defective-RB1-loss (high expressed); (B) OS, DFS, PFS and DSS HR values for different combinations of CD274 (PD-L1 gene) vs (i) RB1 Pathway Activity, Defective-RB1-loss (low expressed) and (ii) Defective-RB1-loss (high expressed); (C) OS, DFS, PFS and DSS HR values for different combinations of AXL gene vs (i) RB1 Pathway Activity, Defective-RB1-loss (low expressed) and (ii) Defective-RB1-loss (high expressed) (* indicates p-value < 0.05, ** indicates p-value <0.01, *** indicates p-value <0.001; OS: Overall Survival, DFS: Disease Free Survival and PFS: Progression Free Survival, DFS: Disease Free Survival)

**Figure S13.**
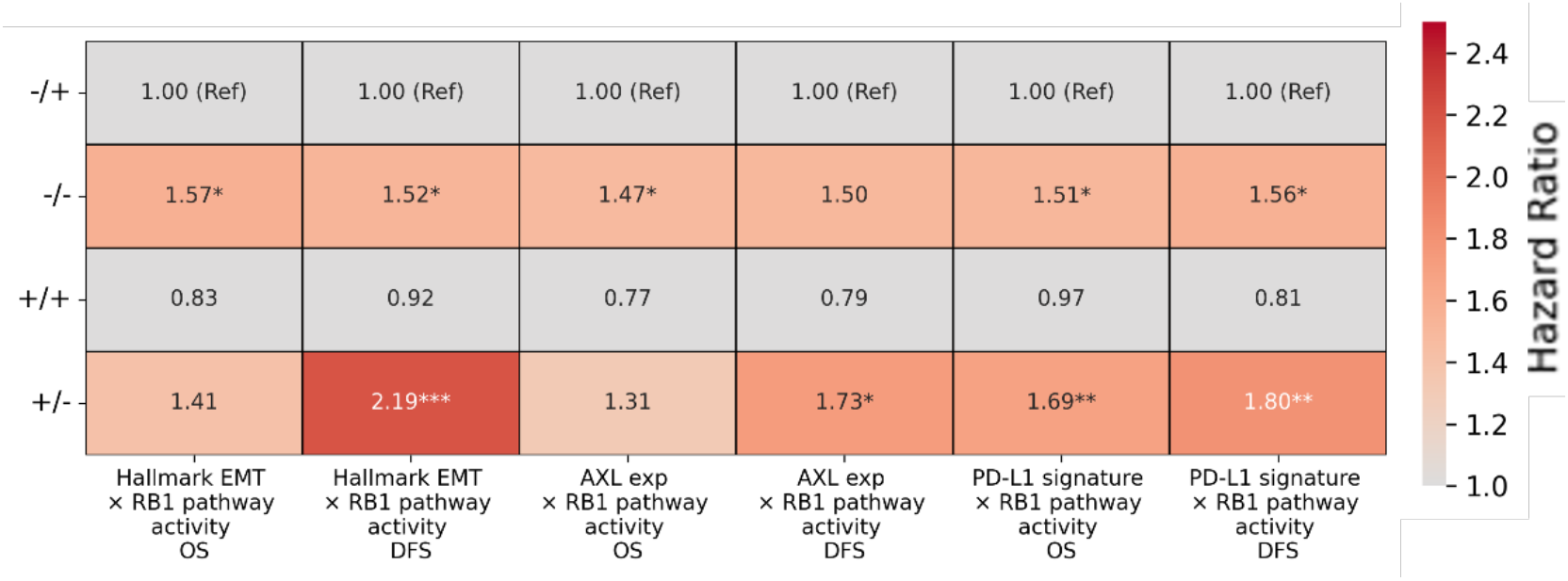
OS and DFS Hazard Ratio (HR) values for double combinations of enrichment scores of different gene-sets and expression levels of genes in GSE68465 patient cohort. OS and DFS HR values for different combinations of Hallmark-EMT vs RB1 Pathway Activity, AXL gene vs RB1 Pathway Activity and PD-L1 Signature vs RB1 Pathway Activity (* indicates p-value < 0.05, ** indicates p-value <0.01, *** indicates p-value <0.001; OS: Overall Survival, DFS: Disease Free Survival)

**Figure S14.**
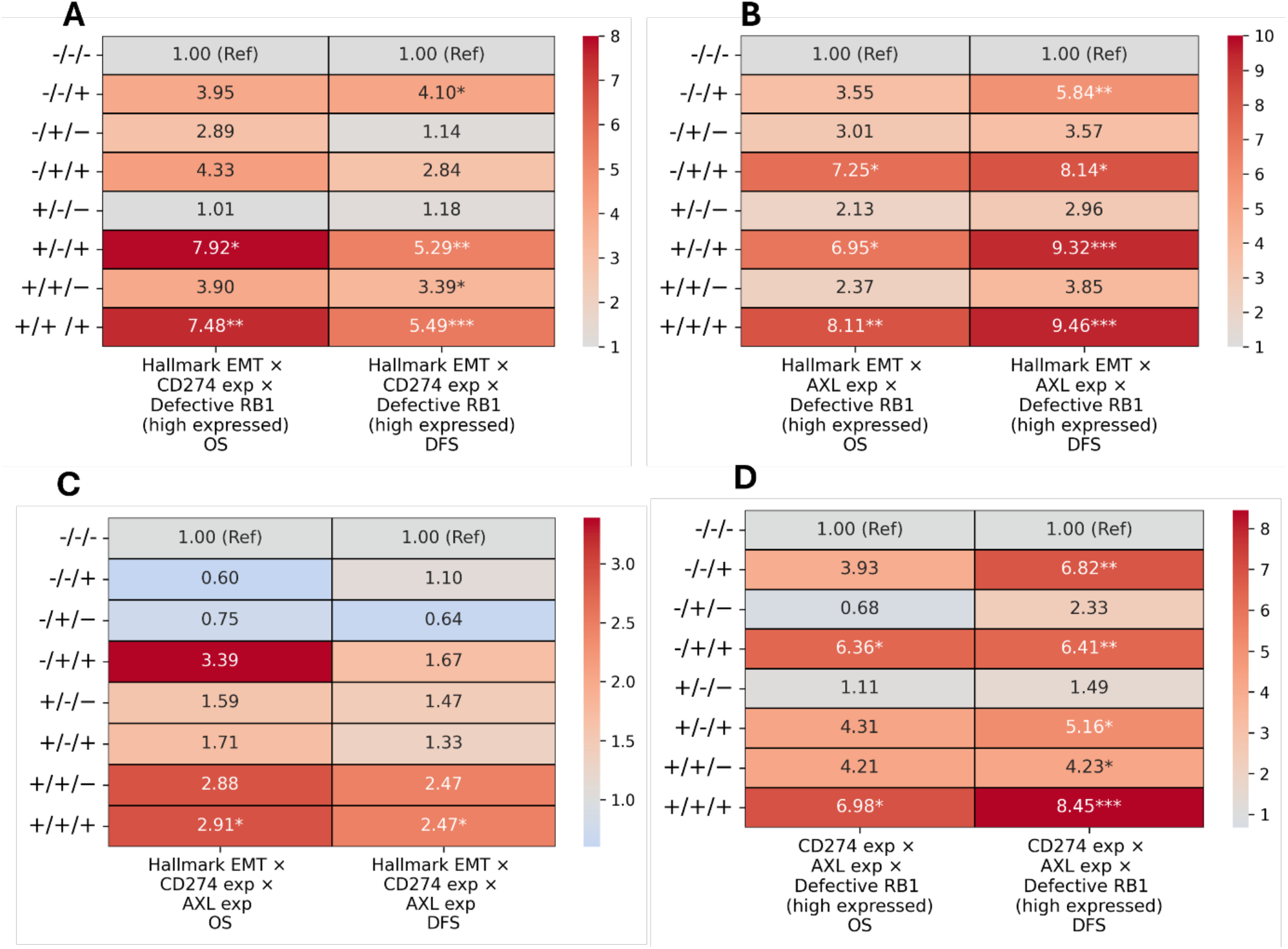
Heatmaps depicting OS and DFS Hazard Ratio (HR) values for triple combinations of enrichment scores of different gene-sets and expression levels of genes in GSE64658 patient cohort. (A) OS and DFS HR values for combination of Hallmark-EMT vs CD274 vs Defective-RB1-loss (high expressed); (B) OS and DFS HR values for combination of Hallmark-EMT vs AXL vs Defective-RB1-loss (high expressed); (C) OS and DFS HR values for combination of Hallmark-EMT vs CD274 vs AXL; (D) OS and DFS HR values for combination CD274 vs AXL vs Defective RB1 (high expressed) (* indicates p-value < 0.05, ** indicates p-value <0.01, *** indicates p-value <0.001; DFS: Disease Free Survival)

**Figure S15.**
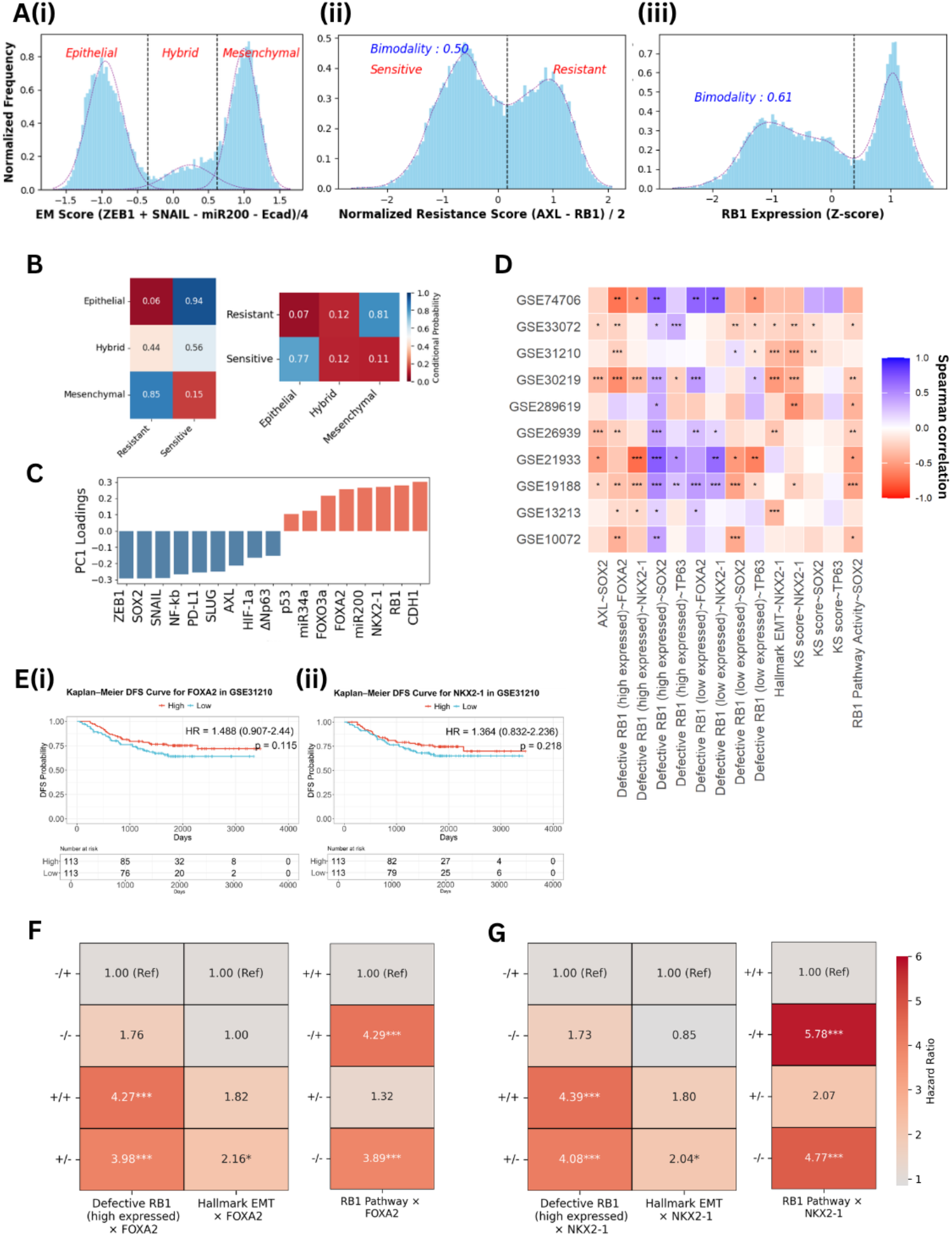
Gene expression and survival analysis for lineage plasticity markers. (A) Density histogram of LUAD-to-LUSC score (= (SOX2 + ΔNp63 – NKX2-1 – FOXA2)/4) fitted to (i) Gaussian mixture model, (i)(ii) Kernel density estimation KDE. (B) Conditional probabilities depicting Epithelial-Sensitive and Mesenchymal-Resistant phenotypes are likely to occur together. (C) Gene steady state values from RACIPE simulations showing LUAD-Epithelial-Sensitive markers having positive PC1 loadings and LUSC-Mesenchymal-Resistant markers having negative PC1 loadings. (D) Correlation values between normalized expression of LUAD maintaining (FOXA2, NKX2-1) and LUSC-switching (SOX-2, TP63) markers with enrichment scores of Hallmark EMT, KS score, AXL and RB1 associated signatures across patient datasets. (E) DFS KM-Plots for gene expression levels of FOXA2 in GSE31210. (F) Heatmaps showing DFS Hazard Ratio (HR) values for different combinations FOXA2 gene expression with RB1-pathway activity, Defective RB1 (high expressed) and Hallmark EMT respectively. (G) Heatmaps showing DFS Hazard Ratio (HR) values for different combinations NKX2-1 gene expression with RB1-pathway activity, Defective RB1 (high expressed) and Hallmark EMT respectively in GSE31210.(* indicates p-value < 0.05, ** indicates p-value <0.01, *** indicates p-value <0.001; OS: Overall Survival, DFS: Disease Free Survival and PFS: Progression Free Survival, DFS: Disease Free Survival)

**Figure S 16.**
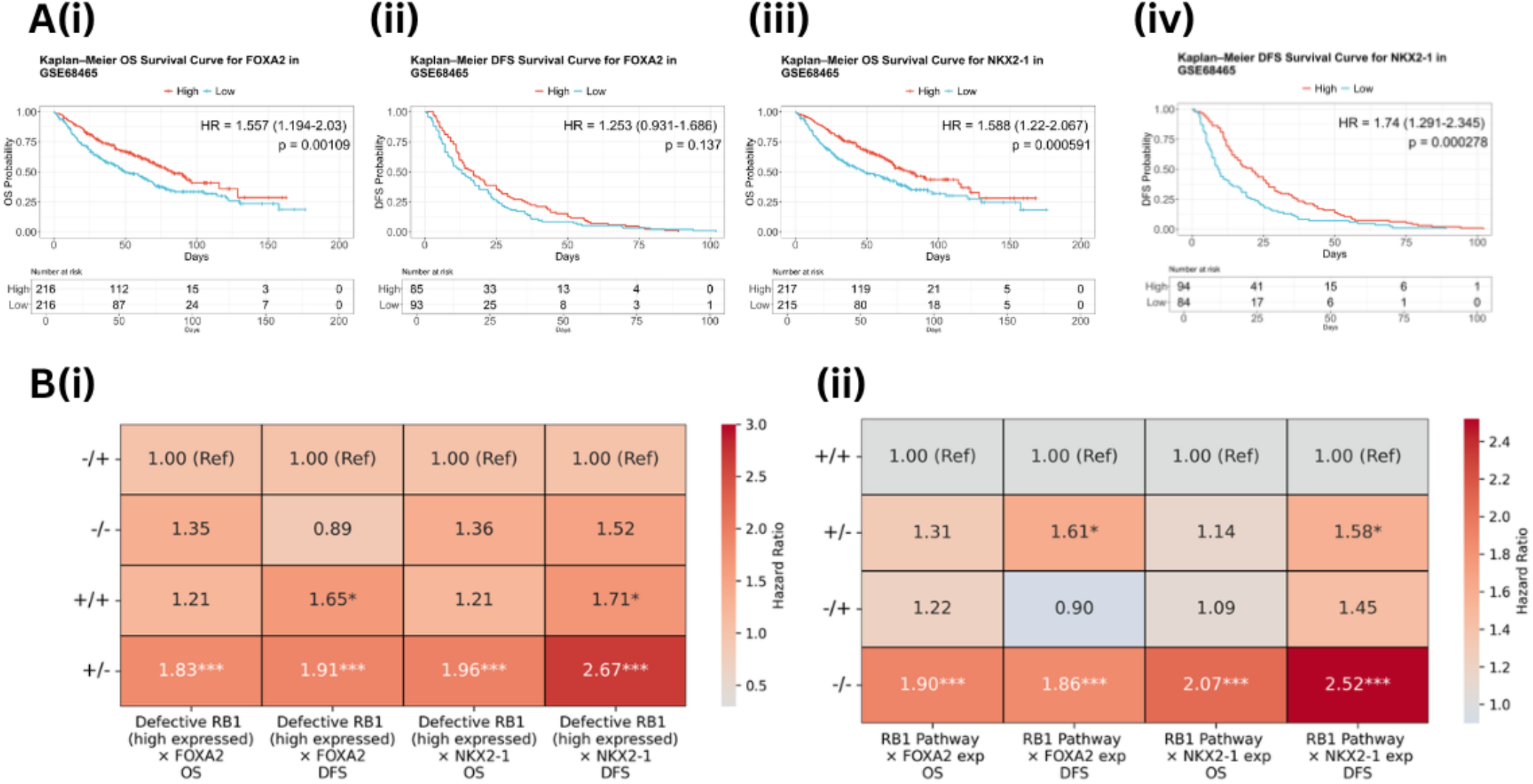
Kaplan Meier and double combination Hazard Ratio (HR) values for OS and DFS for expression levels of NKX2-1 and FOXA2 with enrichment scores of different gene-sets in GSE68465 dataset with patients separated by median. (A)(i, ii, iii, iv) OS and DFS Plots for gene expression levels of FOXA2 and NKX2-1; (B) OS and DFS HR values for different combinations (i) FOXA2 and NKX2-1 gene expression vs Defective-RB1-loss (high expressed) and (ii) ) FOXA2 and NKX2-1 gene expression vs RB1-pathway activity (* p < 0.05, ** p < 0.01, *** p < 0.001; OS: overall survival; DFS: disease-free survival; PFS: progression-free survival)

**Figure S 17.**
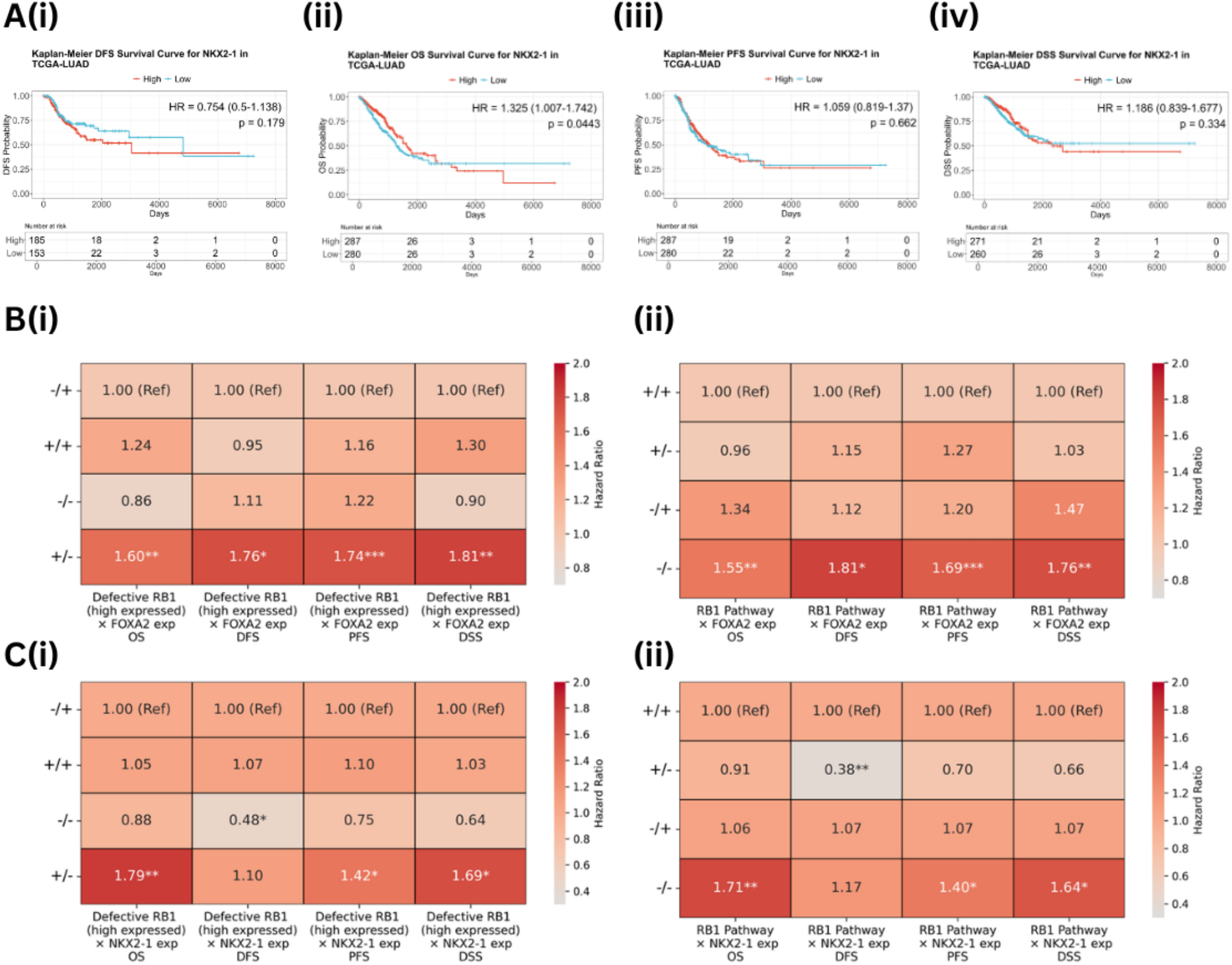
Kaplan Meier and double combination Hazard Ratio (HR) values for PFS, OS, DSS and DFS for expression levels of NKX2-1 and FOXA2 with enrichment scores of different gene-sets in TCGA-LUAD dataset with patients separated by median. (A)(i, ii, iii, iv) OS, DFS, DSS, PFS KM-Plots for gene expression levels of NKX2-1; (B) OS, DFS, DSS and PFS HR values for different combinations (i) FOXA2 gene expression vs Defective-RB1-loss (High expressed) and (ii) FOXA2 gene expression vs RB1-pathway activity; (C) OS, DFS, DSS and PFS HR values for different combinations (i) NKX2-1 gene expression vs Defective-RB1-loss (High expressed) and (ii) NKX2-1 gene expression vs RB1-pathway activity (* p < 0.05, ** p < 0.01, *** p < 0.001; OS: overall survival; DFS: disease-free survival; PFS: progression-free survival)

