## Supplementary Tables for "Bidirectional coupling among EMT, AXL-RB1 signaling and lineage switch drives resistance to osimertinib and worse clinical outcomes in NSCLC"

***Table 1 : References for network edges for Fig 1A and Fig S2***

| S.no | Source | Target | Interaction Type | Reference |
|---|---|---|---|---|
| 1 | AXL | miR34a | Activation | (Cho et al., 2016) |
| 2 | miR34a | AXL | Inhibition | (Cho et al., 2016) |
| 3 | AXL | NF-κB-p65 | Activation | (Asiedu et al., 2014) |
| 4 | NF-κB-p65 | RB1 | Inhibition | (Song et al., 2022) |
| 5 | RB1 | NF-κB-p65 | Inhibition | (Jin et al., 2019) |
| 6 | NF-κB-p65 | p53 | Activation | (H. Wu & Lozano, 1994) |
| 7 | p53 | NF-κB-p65 | Inhibition | (Yang et al., 2015) |
| 8 | p53 | miR200 | Activation | (Tamura et al., 2015) |
| 9 | p53 | miR34a | Activation | (T.-C. Chang et al., 2007) |
| 10 | miR34a | SNAIL | Inhibition | (Kim et al., 2011; Siemens et al., 2011) |
| 11 | SNAIL | miR34a | Inhibition | (Siemens et al., 2011) |
| 12 | NF-κB-p65 | SNAIL | Activation | (Y. Wu & Zhou, 2010) |
| 13 | NF-κB-p65 | ZEB1 | Activation | (Gu et al., 2015) |
| 14 | ZEB1 | p53 | Inhibition | (Larsen et al., 2016) |
| 15 | ZEB1 | miR200 | Inhibition | (Burk et al., 2008) |
| 16 | miR200 | ZEB1 | Inhibition | (Burk et al., 2008) |
| 17 | ZEB1 | Ecad | Inhibition | (Manshouri et al., 2019) |
| 18 | Ecad | ZEB1 | Inhibition | (Park et al., 2008) |
| 19 | NF-κB-p65 | PD-L1 | Activation | (Asgarova et al., 2018) |
| 20 | RB1 | Ecad | Activation | (Arima et al., 2008; Batsché et al., 1998) |
| 21 | miR34a | PD-L1 | Inhibition | (X. Wang et al., 2015) |
| 22 | miR200 | PD-L1 | Inhibition | (El-Daly et al., 2025; Katakura et al., 2020) |
| 23 | RB1 | ZEB1 | Inhibition | (Dean et al., 2015) |
| 24 | ZEB1 | RB1 | Inhibition | (Chen et al., 2017) |
| 25 | PD-L1 | Ecad | Inhibition | (Yu et al., 2020) |
| 26 | miR200 | SNAIL | Inhibition | (Bracken et al., 2015; Zhao et al., 2018) |
| 27 | SNAIL | miR200 | Inhibition | (Bracken et al., 2015; Zhao et al., 2018) |
| 28 | Ecad | SNAIL | Inhibition | (Nagathihalli et al., 2012) |
| 29 | SNAIL | Ecad | Inhibition | (W. Wang et al., 2023) |
| 30 | p53-m | NF-κB-p65 | Activation | (Weisz et al., 2007) |
| 31 | p53-m | AXL | Activation | (Vaughan et al., 2012) |
| 32 | p53-m | ZEB1 | Activation | (Dong et al., 2012) |

*Table 2 : References for network edges for Fig 7A*

| S.No | Source | Target | Interaction Type | Reference |
|---|---|---|---|---|
| 1 | AXL | SOX2 | Activation | (Pan et al., 2025) |
| 2 | FOXA2 | CDH1 | Activation | (Tang et al., 2011) |
| 3 | FOXA2 | NKX2-1 | Activation | (Orstad et al., 2022) |
| 4 | FOXA2 | SLUG | Inhibition | (Tang et al., 2011) |
| 5 | FOXO3a | HIF-1a | Inhibition | (Cheng et al., 2015) |
| 6 | FOXO3a | ΔNp63 | Inhibition | (Hu et al., 2017) |
| 7 | HIF-1a | AXL | Activation | (Rankin et al., 2014) |
| 8 | HIF-1a | SLUG | Activation | (Liu et al., 2018) |
| 9 | HIF-1a | SNAIL | Activation | (Liu et al., 2018) |
| 10 | NF-kb | FOXO3a | Inhibition | (Chiu et al., 2016) |
| 11 | NKX2-1 | CDH1 | Activation | (Saito et al., 2009) |
| 12 | NKX2-1 | SLUG | Inhibition | (Yamaguchi et al., 2013) |
| 13 | NKX2-1 | SNAIL | Inhibition | (Yamaguchi et al., 2013) |
| 14 | NKX2-1 | SOX2 | Inhibition | (Maeda et al., 2012) |
| 15 | NKX2-1 | ΔNp63 | Inhibition | (Goh et al., 2021) |
| 16 | RB1 | SOX2 | Inhibition | (Voigt et al., 2021) |
| 17 | SLUG | CDH1 | Inhibition | (Hung et al., 2019) |
| 18 | SLUG | SNAIL | Inhibition | (Sundararajan et al., 2019) |
| 19 | SLUG | ZEB1 | Activation | (Wels et al., 2011) |
| 20 | SLUG | ΔNp63 | Inhibition | (Herfs et al., 2010) |
| 21 | SNAIL | SLUG | Inhibition | (Sundararajan et al., 2019) |
| 22 | SNAIL | ΔNp63 | Inhibition | (Herfs et al., 2010) |
| 23 | SOX2 | NKX2-1 | Inhibition | (Mollaoglu et al., 2018) |
| 24 | SOX2 | SLUG | Activation | (Z. Chang, 2019) |
| 25 | SOX2 | SNAIL | Activation | (Z. Chang, 2019) |
| 26 | SOX2 | ΔNp63 | Activation | (Ferone et al., 2016) |
| 27 | ZEB1 | FOXA2 | Inhibition | (Fernandes et al., 2021) |
| 28 | ZEB1 | SOX2 | Activation | (Wellner et al., 2009) |
| 29 | ZEB1 | ZEB1 | Activation | (Preca et al., 2017) |
| 30 | ΔNp63 | AXL | Activation | (Dang et al., 2016) |
| 31 | ΔNp63 | SLUG | Activation | (Lambert et al., 2022) |
| 32 | ΔNp63 | ZEB1 | Inhibition | (Tucci et al., 2012) |

***Table 3 : Description of publicly available perturbation in cell line datasets from Gene Expression Omnibus (GEO)***

| S.No | GSE ID | Cell line | Type | Description | Reference |
|---|---|---|---|---|---|
| 1 | GSE49644 | HCC827 | Bulk | TGF-β treatment for three weeks to induce EMT. | (Sun et al., 2014) |
| 2 | GSE81167 | HCC827 | Bulk | RNA-seq from transfected cell lines overexpressing ZEB1 (pcDNA3.1-ZEB1) and control cells (empty pcDNA3.1 vector) | (Zhang et al., 2016) |
| 3 | GSE193258 | HCC827 HCC2935 | Bulk | RNA-seq from cell lines treated with osimertinib for 21 days; 24 h short washout post-treatment and long washout until proliferation resumed | (Criscione et al., 2022) |
| 4 | GSE17708 | A549 | Bulk | Affymetrix microarray expression data from 5 ng/mL TGF-β treatment for 0, 0.5, 1, 2, 4, 8, 16, 24, and 72 h to induce EMT | (Sartor et al., 2010) |
| 5 | GSE14705 | A549 | Single cell | 12 EMT time course with TGF-β treatment | (Cook & Vanderhyden, 2020) |

***Table 4: Description of publicly available patient datasets from Gene Expression Omnibus (GEO)***

| S.No | GSE ID | #patients | Type | Description | Reference |
|---|---|---|---|---|---|
| 1 | GSE31210 | 226 | Bulk | Affymetrix Microarray dataset from 226 lung adenocarcinomas (127 with EGFR mutation, 20 with KRAS mutation, 11 with EML4-ALK fusion and 68 triple negative cases) | (Okayama et al., 2012) |
| 2 | GSE289619 | 32 | Bulk | RNA-seq data from patients with EGFR mutant NSCLC who received osimertinib monotherapy as a first-line treatment between January 2016 and April 2023 | (Ibusuki et al., 2025) |
| 3 | GSE10072 | 58 | Bulk | Affymetrix Microarray dataset from 58 LUAD tumour tissues | (Landi et al., 2008) |
| 4 | GSE11969 | 149 | Bulk | Microarray expression profile for NSCLC tumours | (Matsuyama et al., 2011) |
| 5 | GSE13213 | 117 | Bulk | Microarray expression profiles from LUAD patients | (Tomida et al., 2009) |
| 6 | GSE19188 | 91 | Bulk | Affymetrix Microarray expression data from early-stage NSCLC patients | (Hou et al., 2010) |
| 7 | GSE21933 | 21 | Bulk | Microarray expression profiles from 10 early stage (stages I and II) and 11 late stage (stages III and IV) NSCLC patients | (Lo et al., 2012) |
| 8 | GSE26939 | 116 | Bulk | RNA-seq from LUAD patients | (Wilkerson et al., 2012) |
| 9 | GSE30219 | 293 | Bulk | Affymetrix Microarray expression data from lung tumour samples | (Rousseaux et al., 2013) |
| 10 | GSE33072 | 131 | Bulk | Affymetrix Microarray expression from BATTLE trial with patients treated with first generation EGFR-TKIs | (Criscione et al., 2022) |
| 11 | GSE33532 | 20 | Bulk | Affymetrix microarray expression data for 4 tumor samples each from early-stage NSCLC patients | - |
| 12 | GSE74706 | 18 | Bulk | Affymetrix Microarray expression data from human NSCLC tissues | (Marwitz et al., 2016) |
| 13 | GSE189357 | 9 | Single cell | Droplet-based scRNA-seq from resected samples from nine LUAD patients, three each from adenocarcinoma in situ (AIS, tumour diameter ≤ 3 cm) to minimally invasive adenocarcinoma (MIA, tumor diameter ≤ 3 cm, invasive growth ≤ 5 mm) and invasive adenocarcinoma (IAC, invasive growth > 5mm) | (Zhu et al., 2022) |
| 14 | GSE189487 | 9 | Spatial | Same as (13) GSE189357 | (Zhu et al., 2022) |
| 15 | GSE68465 | 442 | Bulk | Affymetrix Microarray expression data from LUAD patients combined with clinical data used for survival analysis | (Shedden et al., 2008) |
